# Light-induced trapping of endogenous proteins reveals spatiotemporal roles of microtubule and kinesin-1 in dendrite patterning of *Drosophila* sensory neurons

**DOI:** 10.1101/2023.09.30.560303

**Authors:** Yineng Xu, Bei Wang, Inle Bush, Harriet AJ Saunders, Jill Wildonger, Chun Han

## Abstract

Animal development involves numerous molecular events, whose spatiotemporal properties largely determine the biological outcomes. Conventional methods for studying gene function lack the necessary spatiotemporal resolution for precise dissection of developmental mechanisms. Optogenetic approaches are powerful alternatives, but most existing tools rely on exogenous designer proteins that produce narrow outputs and cannot be applied to diverse or endogenous proteins. To address this limitation, we developed OptoTrap, a light-inducible protein trapping system that allows manipulation of endogenous proteins tagged with GFP or split GFP. This system turns on fast and is reversible in minutes or hours. We generated OptoTrap variants optimized for neurons and epithelial cells and demonstrate effective trapping of endogenous proteins of diverse sizes, subcellular locations, and functions. Furthermore, OptoTrap allowed us to instantly disrupt microtubules and inhibit the kinesin-1 motor in specific dendritic branches of *Drosophila* sensory neurons. Using OptoTrap, we obtained direct evidence that microtubules support the growth of highly dynamic dendrites. Similarly, targeted manipulation of Kinesin heavy chain revealed differential spatiotemporal requirements of kinesin-1 in the patterning of low- and high-order dendritic branches, suggesting that different cargos are needed for the growth of these branches. OptoTrap allows for precise manipulation of endogenous proteins in a spatiotemporal manner and thus holds great promise for studying developmental mechanisms in a wide range of cell types and developmental stages.

## INTRODUCTION

Animal development involves a myriad of molecular events that occur at specific times and locations. The same molecular event can result in vastly different outcomes depending on the spatiotemporal properties of the biological context. Spatiotemporal regulation is particularly important for the development of neurons: Spatially, neurons can span broad domains and receive distinct signaling inputs at different compartments because of unique interactions with the surrounding microenvironment. Temporally, neurons undergo distinct stages of differentiation, including neuronal migration, axon pathfinding, dendrite arborization, and synapse formation, before forming functional circuits. Thus, a deeper understanding of the assembly of the nervous system requires effective approaches to probe the spatiotemporal properties of molecular events in neurons.

However, traditional loss-of-function (LOF) approaches, such as gene mutation and RNA interference (RNAi), and gain-of-function (GOF) approaches, such as gene overexpression, are insufficient for extracting fine spatiotemporal information, because they typically affect the entire cell for a long duration. This caveat is particularly limiting for studying proteins that play multiple roles in different parts of the cell or at different stages of differentiation. An example is the microtubule (MT) cytoskeleton, which provides mechanical support to neuronal branches and serves as tracks for motor-based cargo transport (Iwanski and Kapitein, 2023; Kelliher et al., 2019). MT assembly, organization, and dynamics are important for dendrite growth and maintenance (Conde and Caceres, 2009), and MT-based motors, including kinesin and dynein, are important for neuronal polarity and dendrite patterning (Delandre et al., 2016). However, how MTs and MT-based motors control specific aspects of dendrite morphogenesis, such as arbor size and location of dynamic branches, at different stages of neuronal development remains poorly understood. While optogenetic techniques to regulate MTs and motors are being developed (Dema et al., 2023; Liu et al., 2022; Lu et al., 2020; Meiring et al., 2022; Nijenhuis et al., 2020), it remains a challenge to manipulate MTs and their motors in neurons inside animals with high spatiotemporal precision.

In recent years, optogenetics has emerged as a powerful approach for finely dissecting mechanisms of animal development. Taking advantage of protein modules that change confirmations or binding affinities upon activation by light of specific wavelengths, light-controllable agents have been developed to produce specific signaling outputs (Fan and Lin, 2015; Kwon and Heo, 2020; Wu et al., 2011; Zhang and Cui, 2015). These tools offer an unprecedented level of control and specificity, allowing for precise and instantaneous manipulation of molecular events in cells. Despite these benefits, most existing optogenetic tools can only manipulate specific designer proteins and require their overexpression to produce dominant-active effects. Thus, it has been difficult to apply a single optogenetic system to manipulate a wide range of proteins, especially endogenous proteins expressed by animals.

To solve these problems, we developed OptoTrap, a versatile optogenetic system in *Drosophila* that allows *in vivo* manipulation of endogenous proteins that are tagged with GFP or a split GFP fragment. This system acts through light-induced trapping or clustering of the protein of interest (POI) and can be used to probe spatiotemporal roles of the protein in specific cell types. We first characterize the kinetics of this system in epithelial cells and neurons and then demonstrate the effectiveness of protein trapping/clustering in both cell types. To understand how MTs and their associated motors control dendrite patterning, we used OptoTrap to manipulate α-tubulin and kinesin-1 in *Drosophila* class IV dendritic arborization (C4da) neurons, somatosensory neurons that elaborate complex dendritic arbors on the larval epidermis (Grueber et al., 2003). Our investigation reveals the critical role of MTs in maintaining the growth dynamics of terminal dendrites and differential temporal requirements of kinesin-1 in the patterning of low- and high-order dendritic branches.

## RESULTS

### Design of the OptoTrap system

To achieve protein trapping by light, we initially tested the LARIAT design in *Drosophila* sensory neurons. The LARIAT system (Lee et al., 2014) consists of CRY2 and CIB1-MP, where MP is the oligomerization domain of the Calcium/calmodulin-dependent protein kinase IIα (CaMKIIα). CRY2 and CIB act as monomers in dark but bind to each other under blue light (Kennedy et al., 2010). Because CRY2 simultaneously undergoes blue light-dependent oligomerization (Bugaj et al., 2013), the LARIAT system can react to blue light by forming large protein aggregates, which may serve as a base for trapping proteins of interest. We expressed *UAS*-driven mCardinal-CRY2 (mCard-CRY2) and/or CIB-BFP-MP in C4da neurons and examined their responses to blue light. As expected, mCard-CRY2 efficiently diffused into neuronal dendrites and formed blue light-dependent clusters in both thick and fine dendrites (Figure S1A). However, CIB-BFP-MP was mostly retained in the cell body as large aggregates (Figure S1B), possibly related to the large size of its dodecamer (12mer). Unexpectedly, we found that CIB-MP constitutively trapped mCard-CRY2 into aggregates even in the absence of blue light (Figure S1C). We further tested CRY2^D387A^, a light-insensitive mutant of CRY2 that was previously reported not to interact with CIB in yeast and human cells (Lee et al., 2014; Liu et al., 2008), but observed similar aggregation with CIB-MP (Figure S1C), suggesting that the mCard-CRY2/CIB-MP interaction we observed was unlikely due to unintended light exposure. With these results, we concluded that the LARIAT system is ineffective for light manipulation in *Drosophila* neurons.

As an alternative approach for light-inducible protein trapping, we designed an OptoTrap (for Optogenetic Trapping) system that is built upon two blue light-activatable modules, CRY2olig and Magnets. CRY2olig is a CRY2 mutant that exhibits greater ability to cluster upon blue light stimulation (Taslimi et al., 2014). Magnets consist of a positive Magnet (pMag) and a negative Magnet (nMag), both of which exist as monomers in dark but heterodimerize with each other under blue light (Kawano et al., 2015). In OptoTrap, CRY2olig is fused with three copies of pMag, while nMag is fused with a prey protein. Under blue light, CRY2olig-pMag(x3) clusters via the CRY2olig moiety, and nMag-prey is recruited to the cluster via pMag-nMag interaction. By tagging a protein of interest (POI) with a bait protein that constitutively binds to the prey, the POI can be trapped in the aggregates (Figure 1A). In addition to one copy of nMag (1n) fused to the prey, we further developed a variant by fusing the prey to a tandem dimer of nMag (2n). The latter variant is expected to produce larger aggregates due to crosslinking among CRY2olig clusters by pMag-nMag interactions (Figure 1B) and thus is expected to increase the efficiency of trapping.

**Figure 1.**
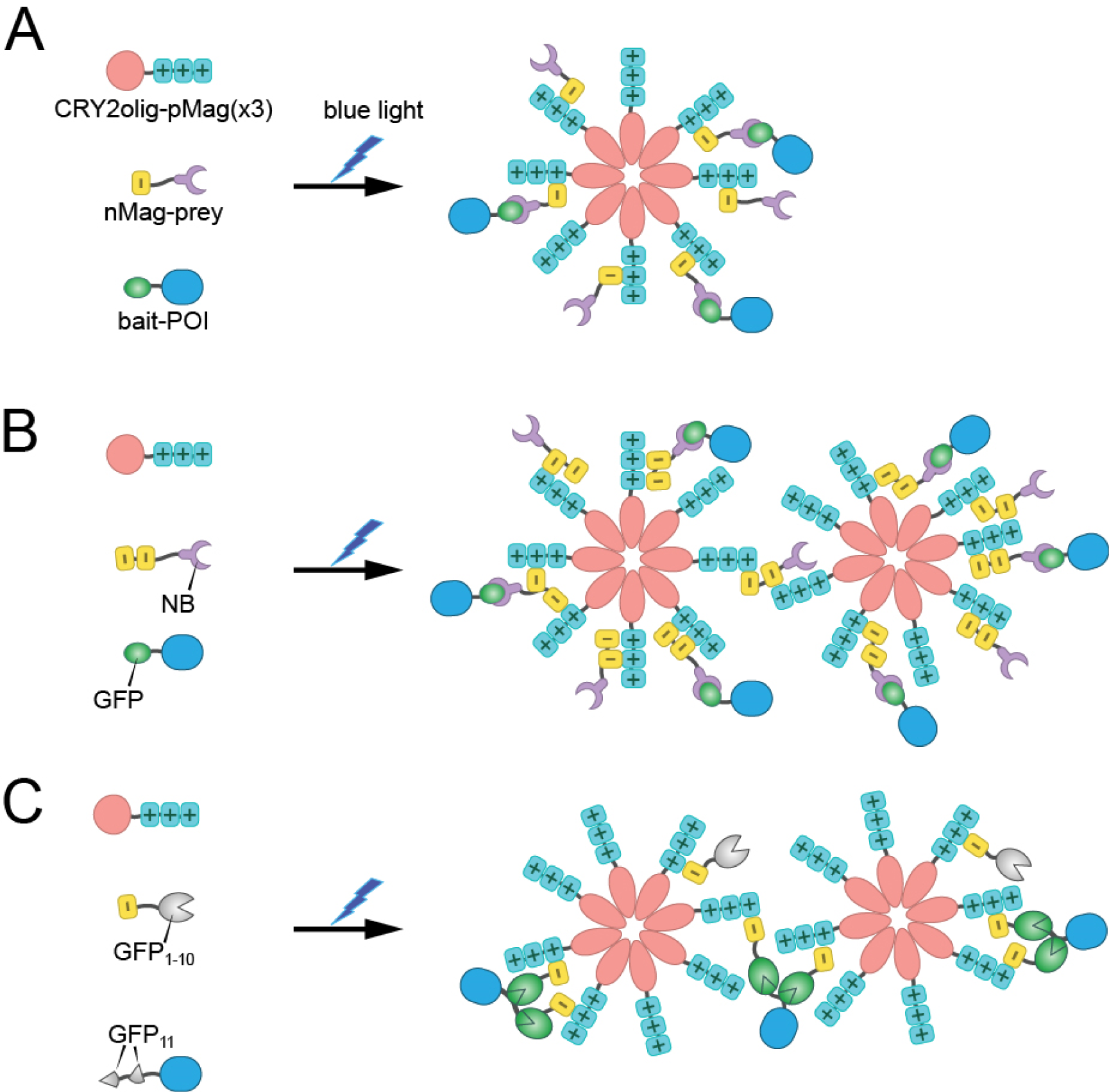
Design of the OptoTrap system. (A) General design of OptoTrap, exemplified by the one nMag (1n) version. POI, protein of interest. (B) OptoTrap with two copies of nMag (2n), nanobody (NB) as the prey, and GFP as the bait. (C) OptoTrap with GFP_1-10_ as the prey and GFP_11_ as the bait.

To increase the versatility of OptoTrap, we utilized two prey-bait pairs: NB-GFP (Figure 1B) and GFP_1-10_-GFP_11_ (Figure 1C). NB is a high-affinity nanobody against GFP (Kirchhofer et al., 2010), while GFP_1-10_ and GFP_11_ are two GFP fragments that automatically reconstitute into a complete fluorescent molecule (Cabantous et al., 2005). By tagging the POI with several tandem copies of GFP_11_, POI is predicted to be trapped into aggregates more efficiently and the aggregates are expected to be bigger due to the crosslinking effect (Figure 1C). In addition, we incorporated two variants of pMag that exhibit different disassociation kinetics: pMag (S) and pMagFast2 (F). The dissociation between pMag and nMag typically takes hours, while pMagFast2 disassociates from nMag within minutes (Kawano et al., 2015).

In total, we generated 6 OptoTrap variants that differ in the pMag-nMag pair, nMag number, and the prey-bait pair (Table 1). These transgenes are controlled by the UAS enhancer (Brand and Perrimon, 1993), so that they can be expressed in a tissue-specific manner in *Drosophila* by Gal4 drivers.

**Table 1.** Variants of OptoTrap.

| ID | CRY2olig-pMag | nMag-Prey |
| --- | --- | --- |
| F1n | Cry2olig-pMagFast2(3x) | mIFP-nMagHigh1-NB |
| S1n | Cry2olig-pMag(3x) | mIFP-nMagHigh1-NB |
| F2n | Cry2olig-pMagFast2(3x) | mIFP-nMagHigh1(2X)-NB |
| S2n | Cry2olig-pMag(3x) | mIFP-nMagHigh1(2X)-NB |
| FG | Cry2olig-pMagFast2(3x) | mIFP-nMagHigh1-GFP<sub>1-10</sub> |
| SG | Cry2olig-pMag(3x) | mIFP-nMagHigh1-GFP<sub>1-10</sub> |

### OptoTrap exhibits light-dependent and reversible aggregation in epithelial cells and neurons

We first assessed the effectiveness of CRY2olig to form light-dependent aggregates in epidermal cells and C4da neurons of *Drosophila* larvae. When using an intermediate epidermal driver (*R15A11-Gal4*), CRY2olig fused to monomeric infrared fluorescent protein (mIFP) appeared diffused in the cell when kept in the dark but formed clusters upon blue laser illumination (Figure 2A). CRY2olig-mIFP exhibited similar light-dependent clustering in dendritic branches of C4da neuron (Figure 2B). The half-times (t_1/2_) of aggregation are 8 s in epidermal cells and 4 s in C4da neurons, as measured by the coefficient of variation (CV) (Figure 2C). Unexpectedly, CRY2olig-mIFP formed aggregates even in the dark when a strong epidermal driver (*R16D01-Gal4*) was used (Figure S2A), suggesting that CRY2olig can form light-independent aggregates at high concentrations in epithelial cells. We did not observe light-independent aggregation of CRY2olig-mIFP in neurons.

**Figure 2.**
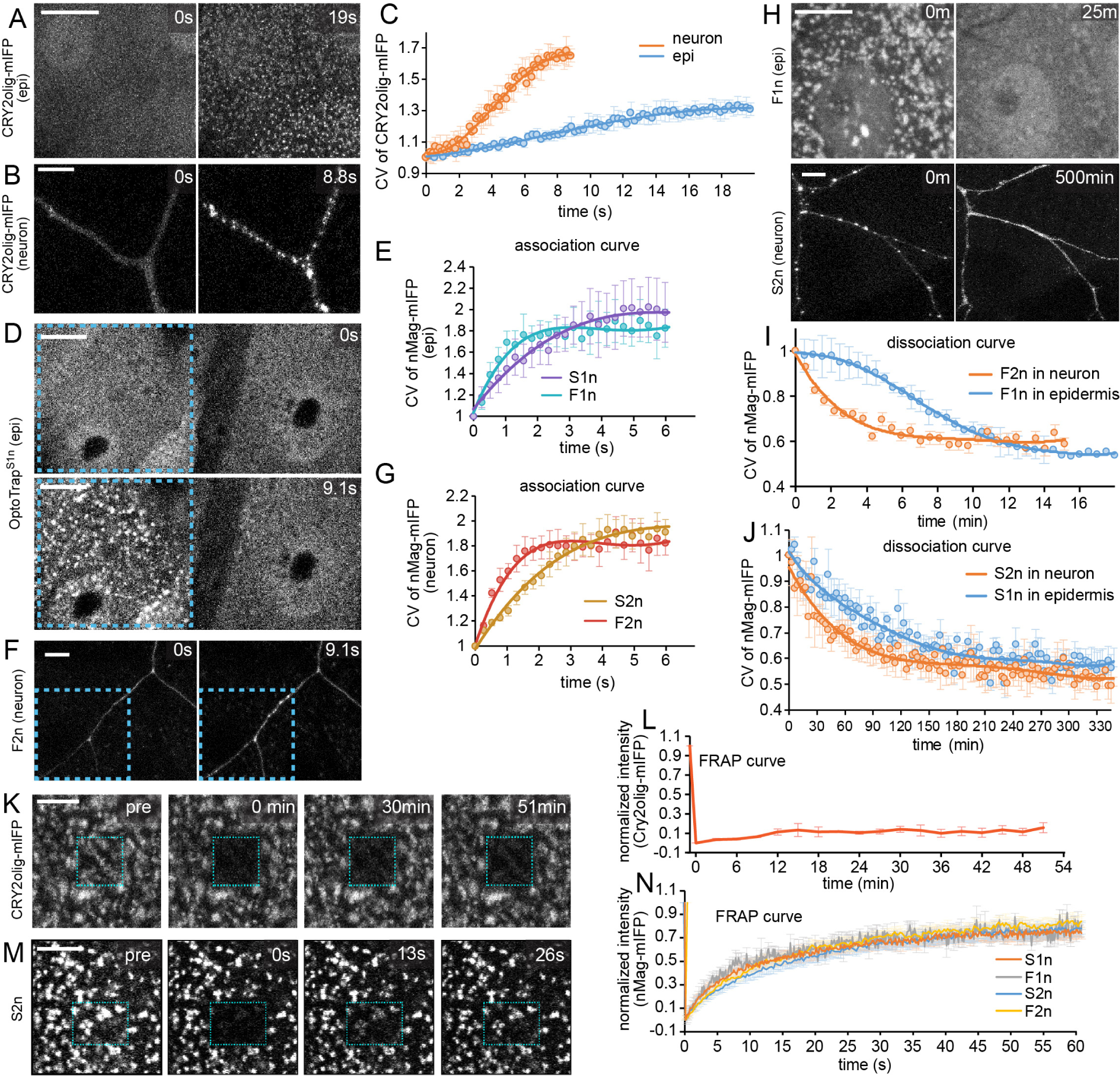
OptoTrap exhibits light-dependent and reversible aggregation in epithelial cells and neurons. (A) Epidermal cells in *Gal4^R15A11^>CRY2olig-mIFP* before (0 s) and after (19 s) blue laser activation. (B) A C4da neuron in *Gal4^ppk^>CRY2olig-mIFP* before (0 s) and after (8.8 s) blue laser activation. *UAS-HO1* was co-expressed in the neuron to make mIFP fluorescent. (C) Coefficient of Variation (CV = standard deviation of mIFP intensity / mean mIFP intensity) of CRY2olig-mIFP in neurons and epidermal cells (epi) plotted over time. n=5 for epi, n=5 for neuron. (D) Epidermal cells in *Gal4^R16D01^>OptoTrap^S1n^* before (0 s) and after (9.1 s) blue laser activation. The illuminated region is enclosed by the dotted line. OptoTrap is visualized by mIFP. (E) CV of OptoTrap mIFP signals in epidermal cells plotted over time. n=9 for S1n, n=7 for F1n. (F) A C4da neuron in *Gal4^ppk^>OptoTrap^F2n^*. The illuminated region is enclosed by the dotted line. *UAS-HO1* was co-expressed to make mIFP fluorescent. (G) CV of OptoTrap mIFP signals in C4da neurons plotted over time. n=5 for S2n, n=6 for F2n. (H) Dissociation of OptoTrap aggregates in epidermal cells (*Gal4^R16D01^>OptoTrap^F1n^*) and C4da neurons (*Gal4^ppk^>OptoTrap^S2n^*). Animals were reared in light but kept in the dark during imaging. (I) CV of mIFP signals in fast versions of OptoTrap in epidermal cells and C4da neurons plotted over time in recovery experiments. n=8 for F1n in epidermis, n=5 for F2n in neuron. (J) CV of mIFP signals in slow versions of OptoTrap in epidermal cells and C4da neurons plotted over time in recovery experiments. n=4 for S1n in epidermis, n=9 for S2n in neuron. (K) Fluorescence recovery after photobleaching (FRAP) of CRY2olig-mIFP in epidermal cells (Gal4^R15A11^>CRY2olig-mIFP). Animals grew in light and were kept in blue laser light during imaging. Blue rectangle indicates the photo-bleached region. (L) Recovery of CRY2olig-mIFP intensity in the bleached region in epidermal cells, normalized by the first frame. n=3 (M) FRAP of OptoTrap mIFP in epidermal cells (*Gal4^R16D01^>OptoTrap^S2n^*). Animals grew in light and were kept in blue light during imaging. Blue rectangle indicates the photo-bleached region. (N) Quantification of OptoTrap mIFP recovery in epidermal cells. n=12 for S1n, n=12 for F1n, n=12 for S2n, n=12 for F2n. All scale bars in image panels represent 10 μm. In plots (C, E, G, I, J), circles, mean; bars, SD; solid line, fit of the curve.

We next evaluated the complete OptoTrap system by measuring the recruitment of mIFP-tagged nMag into aggregates. Interestingly, 1n variants produced more obvious aggregates than 2n variants in epidermal cells upon short illumination (Figures 2D and S2B), while 2n variants are much more effective than 1n variants in neurons (Figures 2F and S2C). Thus, we used 1n only for epidermal cells and 2n only for neurons in subsequent characterization.

In analyses of the switch-on kinetics, aggregation of OptoTrap was rapidly and locally induced in a region of an epidermal cell (Figure 2D; Movie S1) or specific branches of a neuron (Figure 2F; Movie S2), when the blue laser illumination was limited to these regions. The half-times of aggregation are slightly faster for the fast variants (t_1/2_=0.5 s for both F1n in epidermal cells and F2n in neurons) than for the slow variants (t_1/2_=1.2 s for S1n in epidermal cells and t_1/2_=1.6 s for S2n in neurons) (Figures 2E and 2G). We also investigated the switch-off kinetics by monitoring the recovery of nMag-mIFP from aggregates to diffused signals (Figure 2H). The fast variants of OptoTrap had much faster recovery compared to the slow variants: In neurons, F2n showed a t_1/2off_ of 1.6 min, while S2n had a t_1/2off_ of 40 min; in epidermal cells, F1n had a t_1/2off_ of 7.4 min, while S1n had a t_1/2off_ of 65 min (Figures 2I and 2J). Thus, the OptoTrap system contains variants appropriate for investigating biological processes at timescales of minutes to hours in both epithelial cells and neurons.

To better understand protein dynamics within aggregates, we employed Fluorescence Recovery After Photobleaching (FRAP) to measure the diffusion kinetics of CRY2olig and nMag separately in epidermal cells. When kept at the aggregated state, CRY2olig-mIFP showed almost no recovery of signals after photobleaching (Figures 2K and 2L), while nMag-mIFP recovered quickly in all four variants of OptoTrap (t_1/2_=13 s for S1n; 10 s for F1n; 17 s for S2n; 13 s for F2n) to about 80% of the original level (Figure 2M-N). These data suggest that CRY2olig proteins are immobile within aggregates, while most nMag proteins can be exchanged among aggregates. This dynamic property of nMag is compatible with liquid-liquid phase separation and will likely prevent the trapped proteins from forming irreversible aggregates (Zhao et al., 2019).

### OptoTrap can cluster endogenous GFP-tagged proteins under blue light

Taking advantage of the available GFP-tagged endogenous proteins in *Drosophila*, we tested OptoTrap’s versatility in trapping proteins of different sizes and subcellular locations (Table S1) in epidermal cells. These proteins include cytoskeleton-binding proteins, mRNA-binding proteins, enzymes, motor proteins, cell-cell junction proteins. OptoTrap^S1n^ was chosen to trap GFP by NB and to visualize OptoTrap-expressing cells by mIFP fluorescence. *en-Gal4* (Han et al., 2005) was used to express OptoTrap in a strip of epidermal cells in each abdominal segment so that anterior non-expressing cells can serve as an internal control. We employed two light exposure periods: short activation (5 minutes of light after growth in dark) (Figures 3A-3H) and extended activation (120 hours of continuous light throughout the larval development) (Figures 3I-3P). In both methods, we detected OptoTrap-induced aggregates of all proteins in the expression region, which colocalized with OptoTrap. In general, the aggregates induced by short activation were smaller than those by extended activation. Notably, transmembrane junctional proteins Armadillo (Lowe et al.) (Figures 3E and 3M) and Neuroglian (Nrg) (Figures 3H and 3P) were also observed in intracellular puncta colocalizing with OptoTrap. These data suggest that OptoTrap can aggregate and relocate diverse endogenous proteins in a light-dependent manner.

**Figure 3.**
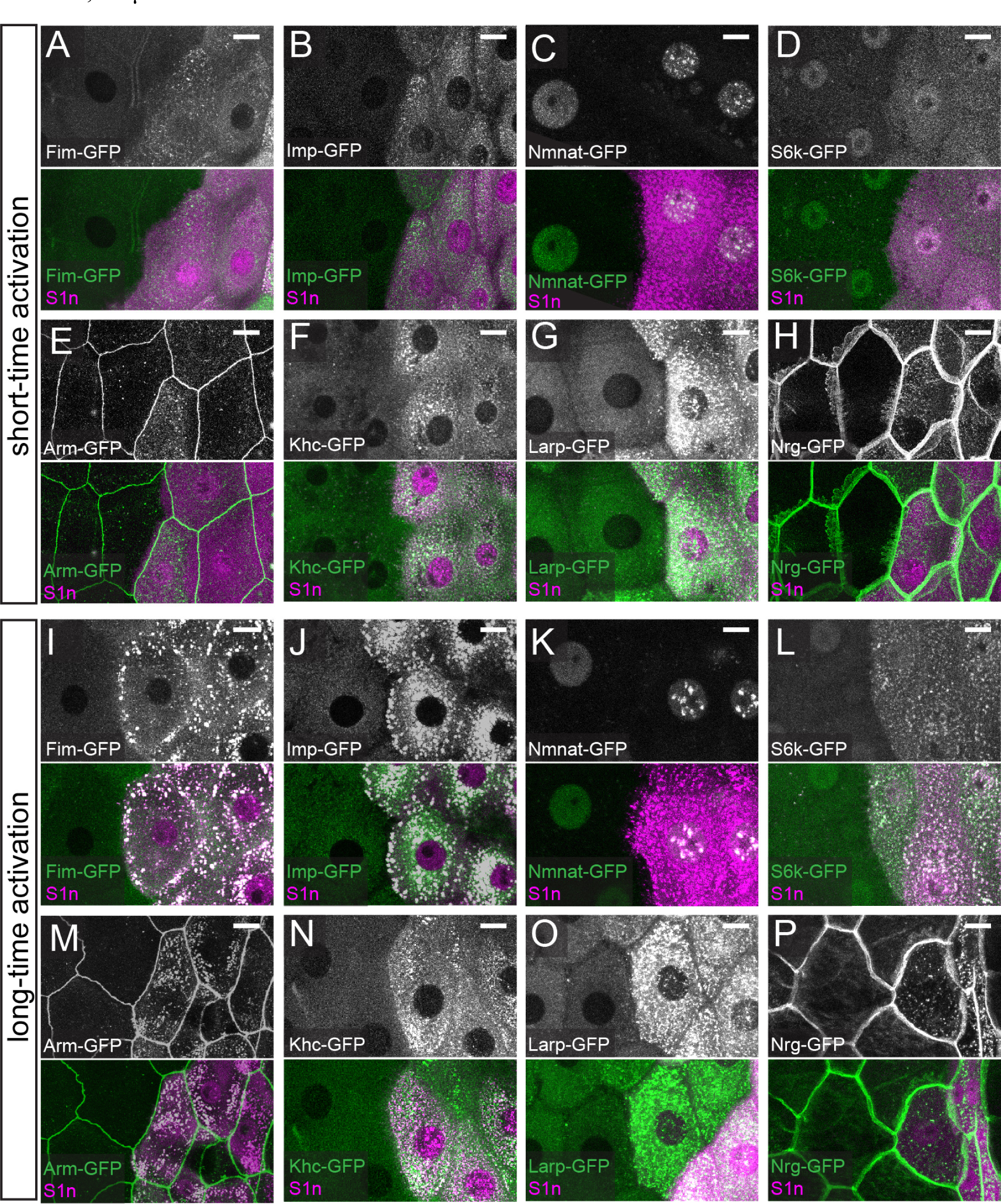
OptoTrap can cluster endogenous GFP-tagged proteins under blue light. (A-H) Various GFP-tagged endogenous proteins in *Gal4^en^>OptoTrap^S1n^*after 5 min blue light illumination. The OptoTrap-expressing epidermal cells are indicated by mIFP (magenta). Scale bars, 25 μm. (I-P) GFP-tagged endogenous proteins in *Gal4^en^>OptoTrap^S1n^* animals that were reared under light since the embryonic stage. Scale bars, 25 μm.

### Trapping of Nrg induces epidermal cell deformation

We next asked whether protein clustering or aggregation induced by OptoTrap can be used to manipulate protein activity by light. Nrg is a component of the septate junction complex (Genova and Fehon, 2003) and is required for the integrity of epithelial septate junctions (Wei et al., 2004). Capable of mediating cell-cell adhesion through homophilic interactions (Hortsch et al., 1995), Nrg has been proposed to activate downstream assembly of cytoskeleton through clustering (Jefford and Dubreuil, 2000). However, no approaches were available previously to directly test this model *in vivo*. We thus examined the impacts of Nrg-GFP clustering by OptoTrap^S1nI^ in a strip of epidermal cells in the middle of every segment. Since the *Nrg* locus is on the X chromosome, we examined both heterozygous females (*Nrg-GFP/+*), in which half of Nrg proteins are tagged by GFP, and hemizygous males (*Nrg-GFP/Y*), in which all Nrg proteins are tagged. Compared to the control in which only mIFP-nMag-NB was expressed (Figure 4A), OptoTrap expression in the dark did not affect epidermal cell shape in either *Nrg-GFP* heterozygotes or hemizygotes (Figures 4B and 4G-4I). However, after 72 hours of blue light exposure, the OptoTrap-expressing cells were deformed in both heterozygotes and hemizygotes (Figure 4C-D). These cells became smaller, as measured by the cell size (Figure 4G), and narrower, as measured by the circularity and the height/width ratio (Figures 4H and 4I). The hemizygotes appeared to have stronger phenotypes than the heterozygotes, but the differences were not statistically significant. In addition, some epidermal cells became multinucleated (arrows in Figures 4C and 4D), possibly due to cell fusion.

**Figure 4.**
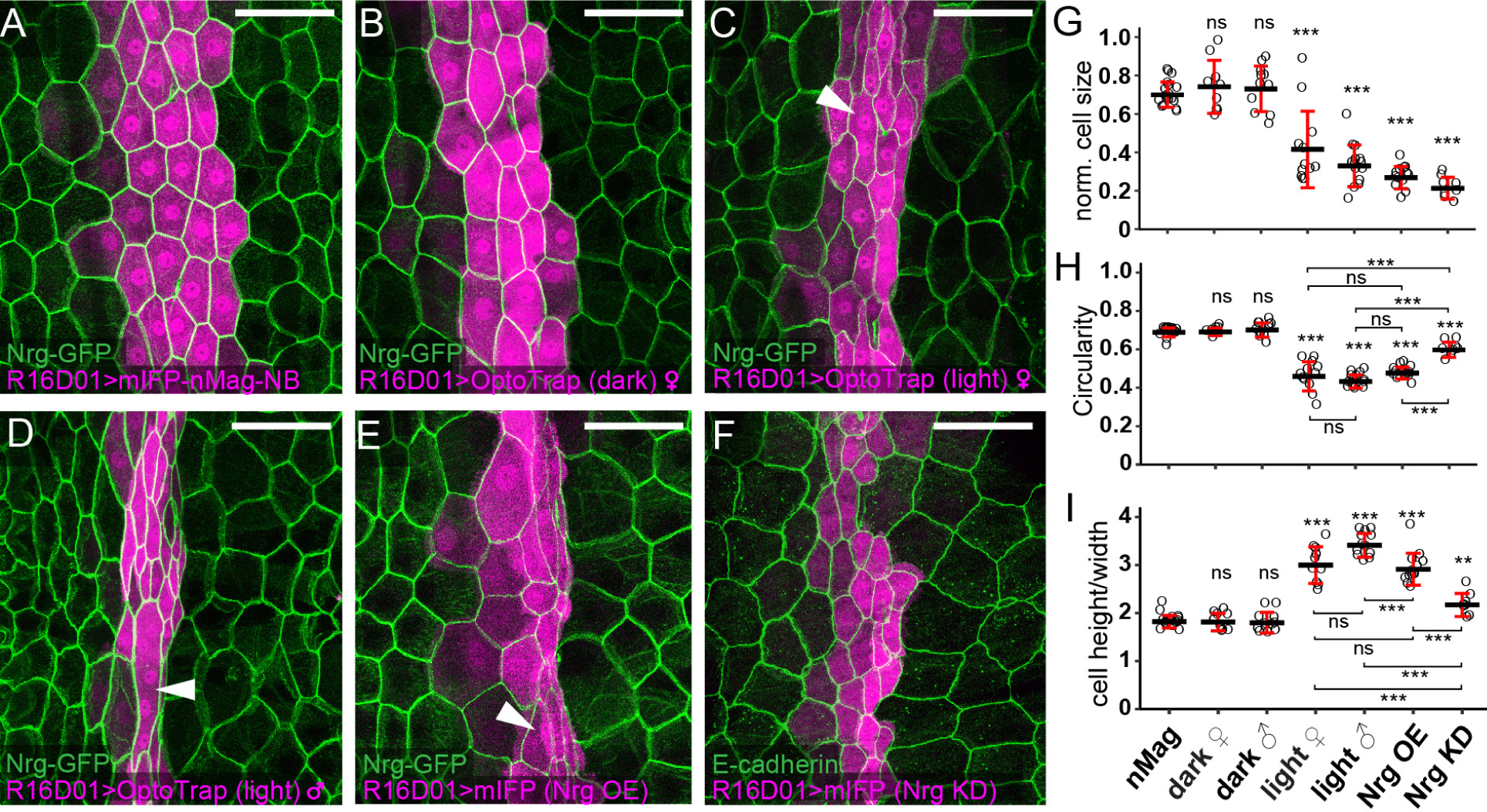
Trapping of Nrg induces epidermal cell deformation. (A-F) Epidermal cells in *Nrg-GFP; Gal4^R16D01^>mIFP-nMag-NB* (A), *Nrg-GFP; Gal4^R16D01^>OptoTrap^S1n^* grown in the dark (B), *Nrg-GFP; Gal4^R16D01^>OptoTrap^S1n^* female grown in light (C), *Nrg-GFP; Gal4^R16D01^>OptoTrap^S1n^* male grown in light (D), *Nrg-GFP; Gal4^R16D01^>UAS-Nrg* (E), and *Gal4^R16D01^>UAS-Nrg-RNAi*. *Gal4^R16D01^*-expressing cells are indicated by mIFP (magenta). The epidermal cell borders are visualized by Nrg-GFP (A-E) and E-cadherin staining (F). Scale bars, 100 μm. (G-I) Normalized size (Gal4^R16D01^ cells / WT cells) (G), Circularity (H), and height/width ratio (I) of *Gal4^R16D01^*-expressing epidermal cells. Each circle represents a segment; n=18 for mIFP-nMag-NB (nMag), n=9 for OptoTrap female (dark), n=11 for OptoTrap male (dark), n=12 for OptoTrap female (light), n=14 for OptoTrap male (light), n=13 for Nrg OE, n=8 for Nrg KD. \*\*\**p*<0.001; \*\**p*<0.01; \**p*<0.05; ns, not significant; One-way ANOVA and Tukey’s HSD test. Black bars, mean; red bars, SD.

The fact that OptoTrap caused epidermal cell deformation in *Nrg-GFP* heterozygotes suggests that the phenotype is dominant, which is consistent with the hypothesis of clustering-induced Nrg activation. We further examined *Nrg* GOF by overexpression (OE) (Wei et al., 2004) and LOF by knockdown (KD). While Nrg KD caused strongest reduction of cell size (Figures 4F and 4G), Nrg OE caused multinucleation and cell narrowing more similarly to OptoTrap manipulations (Figures 4E, and 4G-4I). These data provide supporting, even though not definitive, evidence that clustering induces Nrg activation. More importantly, our results suggest that OptoTrap can be used to manipulate activity of endogenous proteins, including membrane proteins, *in vivo*.

### Optogenetic trapping of α-tubulin results in tunable dendrite reduction of neurons

To investigate the role of MTs in dendrite morphogenesis by OptoTrap, we tagged the *α-Tubulin at 84B* (*αTub84B*) locus with one copy of GFP_11_. Like GFP-tagged *αTub84B* (Jenkins et al., 2017), the *αTub84B-GFP_11_* allele is male sterile; thus, we were unable to obtain *αTub84B-GFP_11_* homozygotes. However, we reason that we may still be able to disrupt MTs with OptoTrap, even if not all the α-tubulins are tagged.

To confirm that αTub84B-GFP_11_ can be aggregated by OptoTrap, we expressed OptoTrap^SG^ in C4da neurons and subjected the larvae to blue light. Expression of nMag-GFP_1-10_ alone resulted in green fluorescence in C4da dendrites (Figure S3A), suggesting successful reconstitution of split GFP. Exposing OptoTrap-expressing larvae to blue light for 5 min led to stronger GFP signals in main dendrite branches, although the signals still appeared to be continuous (Figure S3B). In contrast, with 72 h light exposure, GFP signals were detected in large aggregates in main dendrites (Figure S3C), indicating that OptoTrap^SG^ can trap αTub84B-GFP_11_.

To disrupt MT dynamics, we cultured larvae in the dark for various periods of time, so that GFP_11_-tagged α-tubulin could be assembled into MTs, and then grew the larvae under blue light until they were imaged at 120 h after egg laying (Venken et al.). As controls, neither expressing OptoTrap^SG^ in wildtype (WT) neurons under light nor expressing nMag-GFP_1-10_ only in *αTub84B-GFP_11_* heterozygotes caused dendrite reduction (Figures S3D-S3H), suggesting that OptoTrap aggregation by itself and the reconstitution of GFP on αTub84B do not affect neuronal morphology. As expected, no dendrite reduction was observed in larvae grown in the dark for the entire time (Figures 5A and S3G), indicating the absence of light-independent disruption of MTs. However, in experiments where larvae were switched from dark to light, we observed various degrees of dendrite reduction (Figures 5B-5E, 5G, and 5H). The most severe reduction was observed in a subset of larvae exposed to light for 72 h, where high-order dendrites were nearly absent (Figures 5D, 5G, and 5H). 48 hr light exposure also induced strong dendrite reduction, mainly due to shortening and simplification of high order branches (Figures 5C, 5G, and 5H). Interestingly, animals kept in light for 96 h or longer did not show obvious dendrite reduction (Figure 5E-5H). A possible explanation for this observation is that, with early activation of OptoTrap, most αTub84B-GFP_11_ proteins are sequestered before they could be assembled into MTs, and thus high-order branches mostly contain MTs devoid of tagged α-tubulin.

**Figure 5.**
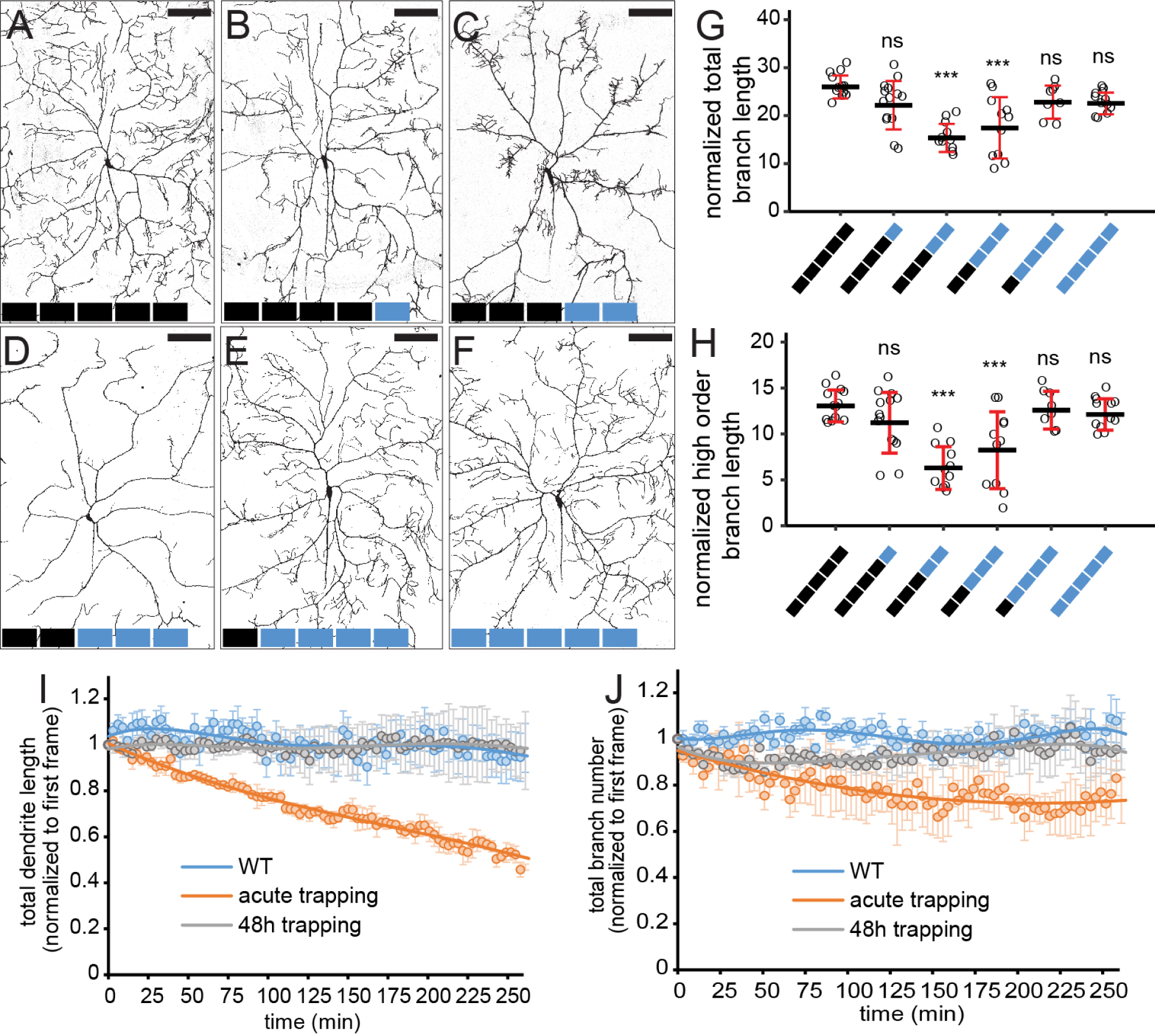
Optogenetic trapping of *α*-tubulin results in tunable dendrite reduction of neurons. (A-F) C4da neurons in *Gal4^ppk^>OptoTrap^SG^; α-Tub84B-GFP_11_/+* animals. Neurons were labeled by *Gal4^ppk^>CD4-tdTom*. Each block represents 24 h either in dark (black) or light (blue). Scale bars, 100 μm. (G and H) Total branch length (G) and high-order (order 5-6) branch length (H) normalized by segment width. Each circle represents a neuron; n=13 for 120 h dark; n=14 for 24 h light and 96 h dark; n=11 for 48 h light and 72 h dark; n=11 for 72 h light and 4 8h light; n=8 for 96 h light and 24 h dark; n=12 for 120 h light. \*\*\**p*<0.001; ns, not significant; One-way ANOVA and Tukey’s HSD test. Black bars, mean; red bars, SD. (I) Total dendrite length of WT neurons (blue line), neurons with α-Tub84B acute trapping (orange line) and 48 h trapping (grey line) plotted over the duration of imaging. The total dendrite length is normalized to that of the first frame. n=8 for WT neuron; n=10 for acute trapping; n=7 for 48 h trapping. Circles, mean; bars, SD; solid line, fit of the curve. (J) Total branch number of WT neuron (blue line), neurons with α-Tub84B acute trapping (orange line) and 48 h trapping (grey line) plotted over the duration of imaging. The total branch number is normalized to that of the first frame. n=8 for WT neuron; n=10 for acute trapping; n=7 for 48 h trapping. Circles, mean; bars, SD; solid line, fit of the curve.

Because MT disruption in the above experiments mainly affected high-order branches, we next used time-lapse imaging (Ji and Han, 2020) to examine dendrite dynamics of three types of neurons: WT neurons, neurons with αTub84B-GFP_11X7_ trapped for 48 h before imaging under light (48 h trapping), and neurons with αTub84B-GFP_11X7_ being trapped from the beginning of the imaging (acute trapping). Within 4.5 h, the total dendrite length and the branch number remained unchanged in WT and 48 h trapping (Figures 5I, 5J, and Movies S3, S4), even though dendrites were reduced in the latter (Figure 5C). These data suggest that neurons had reached a steady state after 48 h trapping. In contrast, acute trapping caused immediate retraction of terminal dendrites (Movie S5) and this trend continued throughout the imaging period (Figures 5I and 5J), indicating that acute tubulin trapping immediately destabilizes terminal dendrites.

We next wondered whether trapping of αTub84B-GFP_11_ can disrupt stable and bundled MTs in thick, low-order dendrites, where MTs form tracks for cargo transport (Zheng et al., 2008). We thus examined MT-mediated cargo transport in these branches using the CD4-tdTomato (CD4-tdTom) marker (Han et al., 2011), which should label all membrane vesicles in the secretory pathway. The majority of CD4-tdTom vesicles in WT dendrites were either static or exhibited retrograde motion, and only a small fraction exhibited anterograde or bidirectional (BD) motion (Figures S3J and S3L). Acute trapping of αTub84B-GFP_11_ did not change this distribution (Figures S3I and S3J). In addition, the distributions of moving vesicle speed were similar in both WT and acutely trapped-tubulin dendrites (Figure S3K). Lastly, we could not detect obvious difference in the distribution of Futsch, a maker of stable and bundled MTs, in thick dendrites after 48 h trapping of αTub84B-GFP_11_ (Figures S3M-S3O). Thus, tubulin trapping does not seem to disrupt stable MTs in thick dendrites, while it can strongly affect dynamic branches.

### Optogenetic trapping causes instant, spatially restricted, and reversible inhibition of kinesin motor in dendrites

MT-based transport plays critical roles in dendrite growth and patterning. The (+) end-directed motor kinesin and the (-) end-directed motor dynein work in concert to transport cargos needed for branch growth to proper locations within the dendritic arbor (Iwanski and Kapitein, 2023; Kelliher et al., 2019). LOF of kinesin and dynein in C4da neurons results in very similar phenotypes of shifting of high-order dendrites towards the cell body (proximal shift) (Satoh et al., 2008; Zheng et al., 2008), suggesting that certain “branching machinery” relies on the motor system for delivery to the distal dendritic arbor. However, how MT-based transport contributes to dendrite growth at different stages of neuronal differentiation remains unknown. To address this question with OptoTrap, we used a *Kinesin heavy chain* (*Khc*) allele tagged with GFP_11x7_ (Kelliher et al., 2018). We reasoned that in *Khc-GFP_11x7_* homozygotes, expressing OptoTrap^FG^ in C4da neurons should allow fast trapping and release of all kinesin-1 motors and thus manipulation of kinesin-mediated transport by light.

When visualized by nMag-GFP_1-10_ only, kinesin-1 is broadly distributed throughout the neuron, including distal terminal dendrites (Figure S4A). This distribution was unchanged by OptoTrap^FG^ expression when animals grew in dark (Figure S4B). OptoTrap caused Khc to form aggregates in these neurons when imaged under blue laser. In comparison, in OptoTrap^FG^-expressing animals subjected to prolonged blue light exposure (72 h), Khc was trapped predominantly in the cell body and sparsely in proximal dendrites, in contrast to its absence in distal dendrites (Figure S4C). These data indicate that OptoTrap efficiently restricts kinesin-1 distribution in neurons in a light-dependent manner.

To test whether Khc trapping is sufficient to inhibit kinesin-1 motor activity, we first examined mitochondrial mobility in dendrites, as mitochondria are cargos of kinesin-1 (Pilling et al., 2006). We applied blue laser to only a portion of the dendrite arbor while imaging mito-mCherry-labeled mitochondria in both illuminated and dark parts of the neuron (Figure 6A). Mitochondria were found to move in both anterograde and retrograde directions in the dark region (Figure 6A’) but appeared static in the illuminated part (Figure 6A’’ and Movie S6). When the entire animal was illuminated, our quantification shows that the percentage of mitochondria exhibiting directional movement reduced from 33% in the WT to 10% in acute trapping and to 1% in chronic (120 h) trapping (Figure 6B). Compared to those in WT neurons, the non-static mitochondria in acute and chronic trapping (nearly all were BD) showed 76% and 85% reduction of speed, respectively (Figure 6C). We next examined the effect of Khc trapping on the transport of CD4-tdTom vesicles, at least a subset of which should be transported by kinesin-1. The percentage of directionally moving CD4-tdTom vesicles reduced from 53% in the WT to 18% in acute trapping and 28% in chronic trapping (Figure 6E). The average speed of moving CD4-tdTom vesicles also reduced 73% in acute trapping and 52% in chronic trapping (Figure 6F). Together, these findings suggest that kinesin-1 can be inhibited instantaneously, locally, and efficiently by OptoTrap, resulting in disruptions of cargo transport.

**Figure 6.**
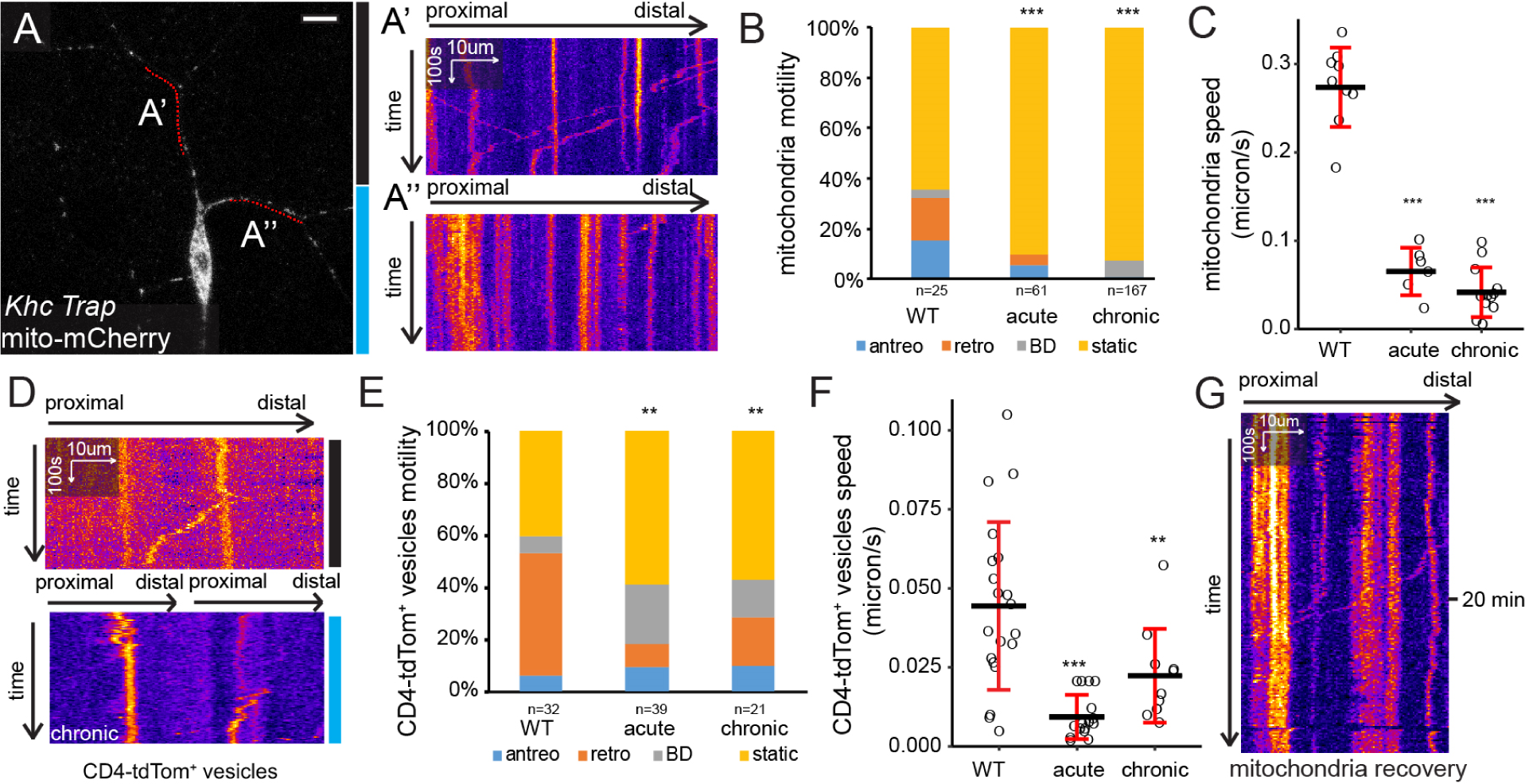
Optogenetic trapping causes instant, spatially restricted, and reversible inhibition of kinesin motor in dendrites. (A-A”) A C4da neuron in *Khc-GFP_11x7_* homozygote that expresses *Gal4^ppk^>OptoTrap^FG^* and the mitochondrial marker mito-mCherry. The animal were reared in dark; the bottom half of the neuron was illuminated by blue laser (blue bar) from the beginning of imaging, and the top half of the neuron remained in the dark (black bar). (A’) and (A”) show kymographs of mitochondria in dendrite branches in the dark (A’) and the illuminated region (A”). Scale bars, 10 μm. (B) Quantification of mitochondria motility. Sample sizes are indicated in the plot. \*\*\**p*<0.001; Freeman–Halton extension of Fisher’s exact test. (C) Speed of CD4-tdTom-labeled vesicles among the moving (anterograde, retrograde and bidirectional motion) populations in (B). Each circle represents a neuron. n= 9 for WT, n=6 for acute, n=12 for chronic. \*\*\**p*<0.001; One-way ANOVA and Tukey’s HSD test. Black bars, mean; red bars, SD. (D) Kymographs of CD4-tdTom-labeled vesicles in C4da neurons of Khc-GFP_11x7_ homozygote expressing OptoTrap^FG^ that were kept in the dark (top) or reared and imaged in light (bottom). (E) Motility of CD4-tdTom-labeled vesicles. Sample sizes are indicated in the plot. \*\**p*<0.01; Freeman–Halton extension of Fisher’s exact test. (F) Speed of CD4-tdTom-labeled vesicles among the moving (anterograde, retrograde and bidirectional motion) populations in (E). Each circle represents a vesicle. n=20 for WT; n=16 for instant; n=10 for chronic. \*\**p*<0.01; One-way ANOVA and Tukey’s HSD test. Black bars, mean; red bars, SD. (G) A kymograph of mitochondria in Khc-GFP_11x7_ homozygote expressing OptoTrap^FG^ that grew in light and kept in the dark while imaging.

Lastly, we investigated whether the inhibition of kinesin-1 by OptoTrap^FG^ is reversible by examining the recovery of mitochondrial mobility. We grew larvae under blue light for 120 h to keep Khc trapped before imaging mitochondria in the dark. As expected, mitochondria were static in the beginning, but some mitochondria began to move within 20 min of imaging (Figure 6G). These results suggest that Kinesin-1 can be released from OptoTrap aggregates and become functional again, even after prolonged trapping.

### Optogenetic trapping reveals temporal and spatial contributions of Khc to dendrite morphogenesis

Having established the effectiveness of OptoTrap in manipulating kinesin-1 activity, we next asked how inhibiting kinesin-1 at different temporal windows may affect the final dendrite pattern. To understand the impacts on the growth of low-order (or primary) v.s. high-order dendrites and proximal-distal distribution of high-order branches, we measured dendrite arbor size, total branch length, high-order (orders 5-6) branch length, and radial distribution of high-order branches (from the soma). As negative controls, we first examined OptoTrap^FG^ expression in the WT under light, nMag-GFP_1-10_ expression alone in *Khc-GFP_11x7_* homozygotes, and OptoTrap^FG^ expression in *Khc-GFP_11x7_* homozygotes that were kept in the dark. *Khc-GFP_11x7_* homozygotes in these experiments were derived from *Khc-GFP_11x7_* homozygous mother to eliminate possible maternal contribution of untagged Khc proteins from the mother’s germline, such that all Khc proteins in these animals are tagged (Figure S5G). Like OptoTrap^SG^ (Figures S3E, S3G, S3H), OptoTrap^FG^ expression in WT neurons did not affect dendrite length or arbor size, except for slightly shifting the radial distribution of high-order branches distally (Figures S5A, and S5D-S5F). Expressing nMag-GFP_1-10_ alone in *Khc-GFP_11x7_*, which is expected to make the kinesin-1 motor bulkier, caused a mild dendrite reduction as reflected by the dendrite arbor size (13% reduction) and the total dendrite length (26% reduction) (Figures S5B-S5F). Interestingly, expressing the complete OptoTrap^FG^ in *Khc-GFP_11x7_* homozygotes, when grown in the dark, resulted in a much weaker (17%) dendritic reduction and no changes in arbor size, high-order branch length or distribution (Figures 7A, S5C-S5F). Thus, we concluded that these minimal impacts on dendrite morphology can serve as a baseline for optogenetic manipulations of Khc. Next, as a positive control where Khc is inhibited to the fullest extent, we grew animals of the same genotype under light for the entire embryonic and larval period (120 h). C4da neurons in these animals exhibited 62% reduction of arbor size, 67% reduction of total branch length, 52% reduction and strong proximal shift of high-order branches (Figures 7F and 7H-K). These severe defects are consistent with the reported phenotype of *Khc* mutant neurons (Satoh et al., 2008), again demonstrating the effectiveness of OptoTrap in inhibiting kinesin-1 in neurons.

**Figure 7.**
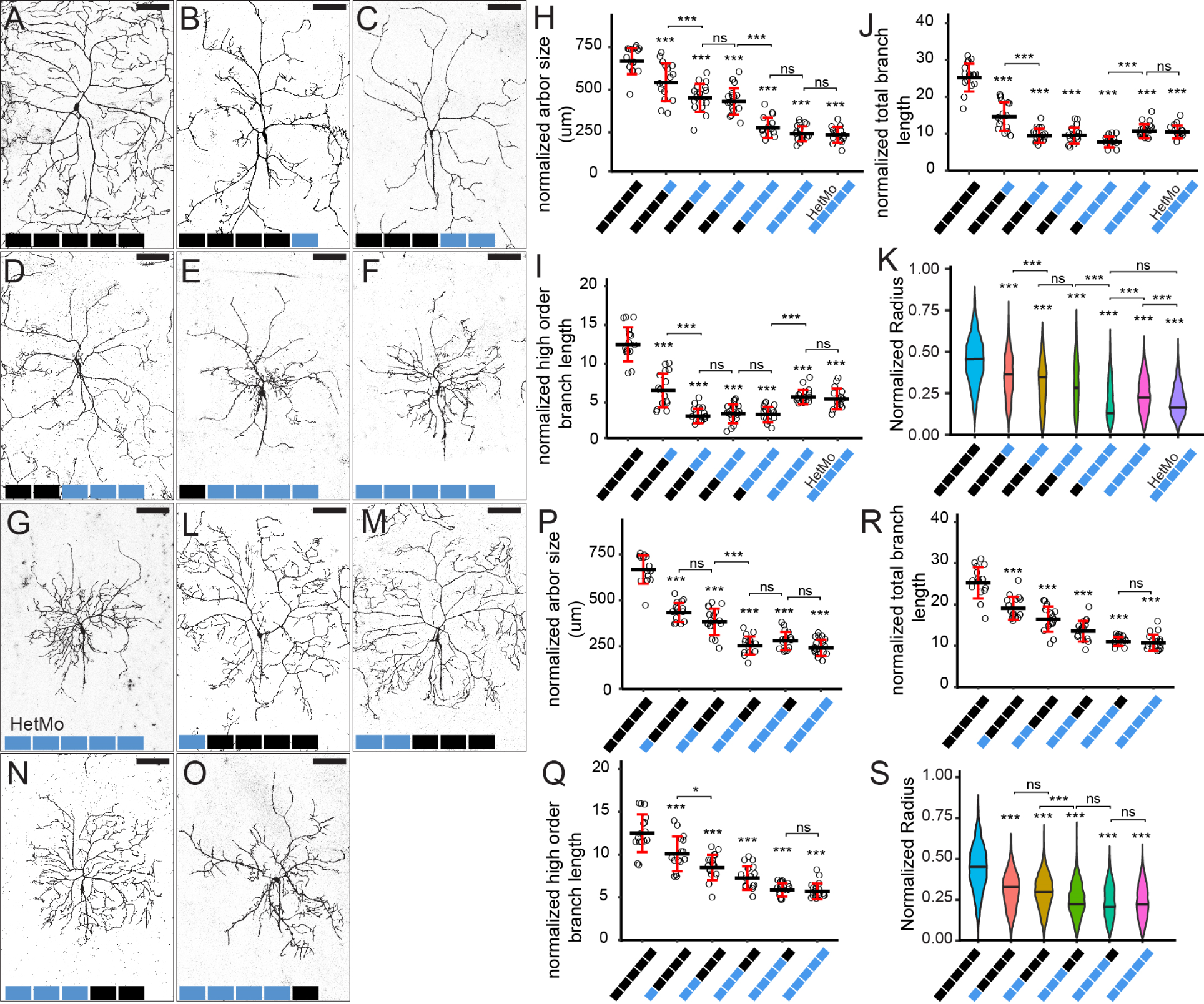
Optogenetic trapping reveals temporal and spatial contributions of Khc to dendrite morphogenesis. (A-G) C4da neurons in *Khc-GFP_11x7_; Gal4^RluA1^>OptoTrap^FG^*animals. Each block represents 24 h either in dark (black) or light (blue). Neurons were labeled by *ppk-CD4-tdTom*. Scale bars, 100 μm. (H-J) Arbor size (H), high-order branch length (I), and total branch length (J) of C4da neurons normalized by segment width. Each circle represents a neuron. (K) Sholl analysis of high-order branches of C4da neurons presented as violin plots. Radius is measured from the cell body and normalized by segment width. The width of violins correlates with the number of intersections between concentric circles with dendrites at a given radius. In (H-K), n=16 for 120 h dark; n=16 for 24 h light and 96 h dark; n=17 for 48 h light and 72 h dark; n=18 for 72 h light and 48 h light; n=15 for 96 h light and 24 h dark; n=18 for 120 h light; n=15 for 120 h light with heterozygous mother (HetMo).. (L-O) C4da neurons in *Khc-GFP_11x7_; Gal4^RluA1^>OptoTrap^FG^* animals. Each block represents 24 h either in dark (black) or light (blue). Neurons were labeled by *ppk-CD4-tdTom*. Scale bars, 100 μm. (P-R) Arbor size (P), high-order branch length (Zhai et al.), and total branch length ® of C4da neurons normalized by segment width. Each circle represents a neuron. (S) Sholl analysis of high-order branches of C4da neurons presented as violin plots. In (P-S), n=16 for 120 h dark; n=15 for 24 h light and 96 h dark; n=17 for 48 h light and 72 h dark; n=16 for 72 h light and 48 h light; n=16 for 96 h light and 24 h dark; n=18 for 120 h light. The 120 h dark and 120 h light datasets are the same as in (H-K). For all quantifications, \*\*\**p*<0.001; \**p*<0.05; ns, not significant; One-way ANOVA and Tukey’s HSD test. Black bars, mean; red bars, SD.

Next, to understand the temporal requirements of kinesin-1 in dendrite patterning, we reared animals first in the dark for various durations and then kept them under light until they were imaged at 120 h AEL (dark-to-light experiments). The neurons were compared to those of the baseline (0 h) and the positive control (120 h). Interestingly, we observed differential effects on low- and high-order dendrites. First, increasing length of Khc inhibition was associated with gradual reduction of the arbor size (Figures 7A-7H), which is primarily determined by the lengths of low-order (1-3) branches. This suggests that kinesin-1 supports the growth of low-order branches throughout animal development. Second, surprisingly, 48 h, 72 h, and 96 h of light exposure resulted in similarly extreme dendrite reduction (69%-73%) (Figure 7J), primarily due to similarly severe reductions of high-order branches (70%-73%) in these groups (Figure 7I). Third, although all groups exposed to light showed reduction of high-order branches, they exhibited distinct high-order branch distributions: While 24 h and 48 h groups showed uniform reductions throughout the arbor (Figures 7B, 7C and 7K), high-order branches in the 96 h group were clustered near the soma and depleted at the distal arbor (Figures 7E and 7K); the 72 h group showed an intermediate phenotype (Figures 7D and 7K). These data suggest that early (24-48 h AEL) kinesin-1 activity is necessary for promoting high-order branch growth at the distal arbor, while kinesin-1activity at later larval development is important for maintaining high-order branches everywhere.

Unexpectedly, we found that 96 h inhibition caused stronger reduction and proximal shift of high-order branches than 120 h (Figures 7E, 7F, 7I and 7K), even though the latter should induce additional Khc inhibition during the first 24 h of animal development. To understand the impact of low Khc activity during early neuronal development, we examined *Khc-GFP_11x7_* homozygotes derived from *Khc-GFP_11x7_/+* heterozygous mothers (HetMo). The neurons in these animals may inherit a small amount of untagged *Khc* mRNA or protein that was deposited maternally into the oocyte (Figure S5G). When grown under light for the whole time, interestingly, these neurons also showed stronger proximal shift than 120 h (Figures 7G and 7K), even though both groups showed similar reduction of arbor size and dendrite length (Figures 7H-7I). These results suggest that maternally contributed Khc can indeed be passed into postmitotic neurons and that residual Khc activity in early neuron development enhances proximal shift and reduction of high-order dendrites.

Next, we asked whether early Khc inhibition has long-lasting effects on the final dendrite pattern or, in other words, whether reactivating Khc later in development can rescue the dendrite defects caused by earlier inactivation. To achieve this, we grew larvae under light for various durations before transferring them to the dark and finally imaging them at 120 h AEL (light-to-dark experiments). Like in dark-to-light experiments, we observed differential effects on low- and high-order dendrites. First, inhibiting Khc in the first three days produced strong effects on the final arbor size: 24 h, 48 h, and 72 h early Khc inhibition resulted in 33%, 41%, and 61% reduction of the arbor size, respectively (Figures 7L-7N and 7P), which are much higher than the 16%, 30%, and 33% reduction of the arbor size caused by late Khc inhibition for the same lengths (Figure 7H). Also, further shortening the recovery time below 48 h did not produce smaller arbors (Figures 7O and 7P). These data suggest that the first three days of animal development are the most critical window for arbor growth and that kinesin-1 activity in the last two days cannot revert the impacts on the arbor size caused by early inhibition. Second, with increasing lengths of early Khc inhibition (0-96 h), we observed increasing degrees of reduction of both total (24%-56%) and high-order (17%-58%) branches (Figures 7R and 7Q). However, the dendritic reductions were much weaker than those in dark-to-light experiments with the same durations of Khc inhibition (Figures 7J and 7I). These data suggest that although the growth of high-order branches is cumulative, Khc activity during later neuronal development is more important than early activity in promoting high-order branch growth. Interestingly, with early suppression, relieving kinesin-1 activity for the last 24 h did not influence the length of total and high-order branches (Figures 7O, 7R, and 7Q). This suggests that it takes longer than 24 h for kinesin-1 to recover the support for high-order branch growth. Lastly, we found that with recovery time longer than 24 h, high-order branches grew uniformly throughout the arbor (Figures 7L-7N and 7S).

Together, the results from our temporal manipulations of Khc suggest that kinesin-1 activity contributes to the growth of low- and high-order branches differentially in temporal-specific and spatial-specific manners. First, while kinesin-1 promotes the arbor growth during the entire development of the neuron, the first 72 h are the most critical window that determines the arbor size. Reintroducing Khc activity after this period has little effect on the arbor size. Second, to cause proximal shift of high-order branches, both persistent suppression of Khc in the last 96 h and early activity within the first 24 h of animal development are necessary. Third, kinesin-1 activity in the last 48 h is necessary for maintaining high-order branches throughout the arbor, but it takes longer than 24 h for reactivated kinesin-1 to regrow high-order branches.

### Optogenetic trapping of Khc disrupts dendrite dynamics

To understand how short-term kinesin-1 inhibition leads to defects of dendrite growth, we examined the dendrite dynamics of C4da neurons in which Khc was acutely inhibited or had been trapped for 24 h. As expected, WT control neurons did not show net changes in total dendrite length or branch number within 70 min (Figure 8A and 8B, blue line). In contrast, both acute inhibition (Figure 8A and 8B, orange line) and 24 h inhibition (Figure 8A and 8B, grey line) groups showed gradual and continuous reductions of both total dendrite length and branch number while being imaged under light, with the latter showing higher rates of reduction. These results are consistent with the graded reduction of dendrites caused by increasing durations of Khc inhibition in the dark-to-light experiments (Figures 7B and 7C) and indicate that the retraction of still-dynamic branches speeds up with longer kinesin-1 inhibition, possibly due to increasing local depletion of growth-promoting factors.

**Figure 8.**
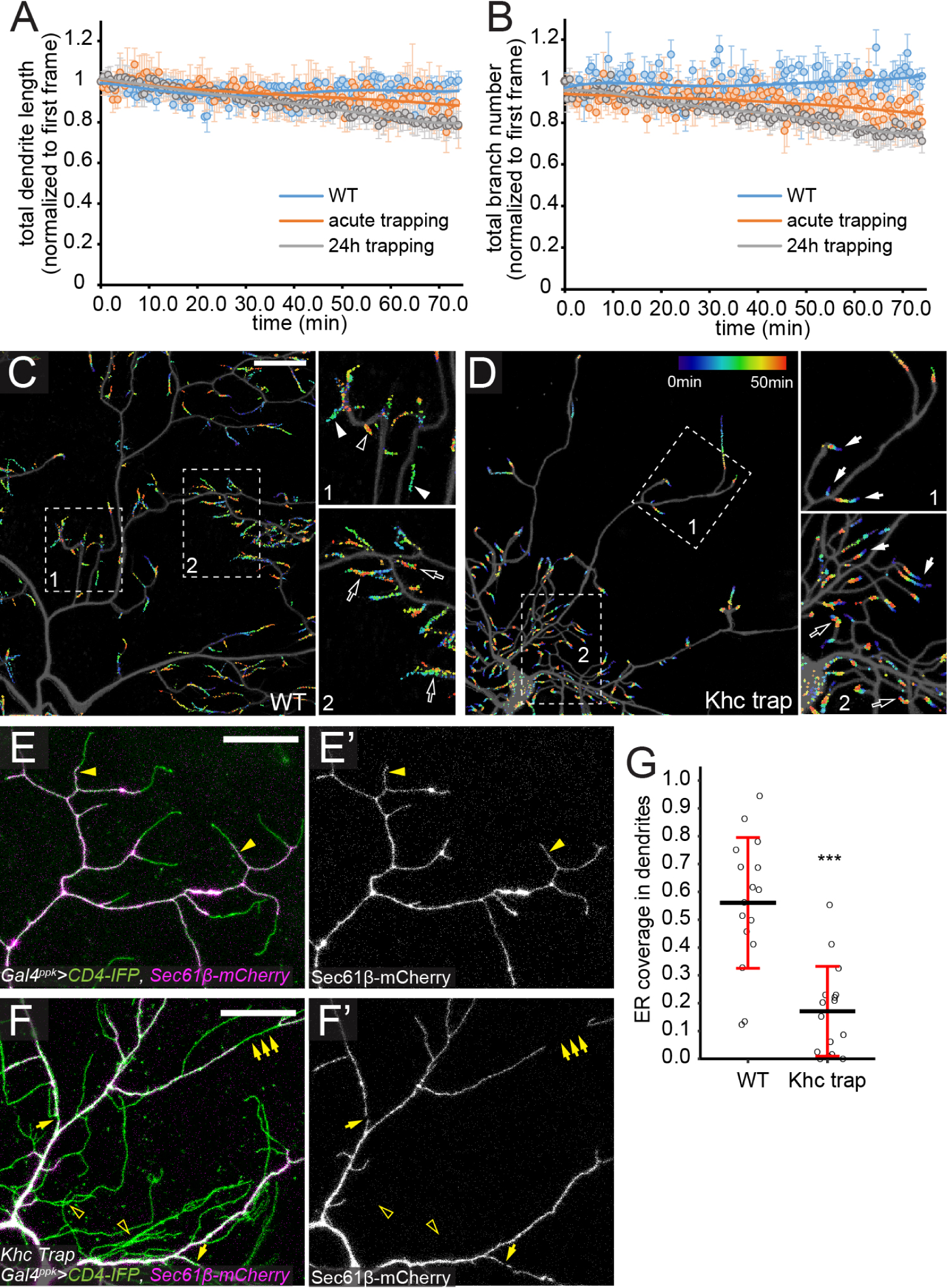
Optogenetic trapping of Khc disrupts dendrite dynamics and ER delivery to terminal dendrites. (A and B) Total dendrite length (A) and total branch number (B) of WT neurons (blue), Khc acute trapping neurons (orange), and Khc 24h trapping neurons (grey) plotted over time. The data were normalized to those of the first frame. n=10 for WT neuron; n=9 for Khc acute trapping neuron; n=9 for Khc 24h trapping neuron. Bars, SD (C and D) Temporal projections of C4da neurons in WT (C) and *Khc-GFP_11x7_; Gal4^ppk^>OptoTrap^SG^* HetMo reared in constant light (D). The positions of dendrite endings in each frame were marked by a color that corresponds to the temporal sequence of the frame (early, blue; late, red). Open arrowheads indicate newly added branches; closed arrowheads indicate eliminated branches; open arrows indicate branches that exhibited repeated retraction and extension; closed arrows indicated branches that slowly retracted. Scale bars, 25 μm. (E-F’) ER distribution in WT (E) and *Khc-GFP_11x7_; Gal4^ppk^>OptoTrap^SG^*HetMo that were reared in light (Khc Trap) (F). ER was marked by *Gal4^ppk^>Sec61β-mCherry* and dendrites were labeled by CD4-IFP. Closed yellow arrowheads indicate ER in terminal branches; open yellow arrow heads indicate the lack of ER in high-order branches; yellow arrows indicate ER breakage in Khc Trap neurons. Scale bars, 25 μm. (H) ER coverage in high-order branches in WT and Khc trap neurons. Each circle represents a neuron; n=16 for WT; n=16 for Khc trap \*\*\**p*<0.001; One-way ANOVA and Tukey’s HSD test. Black bars, mean; red bars, SD.

Next, to understand how prolonged Khc inhibition affects the kinetics of dendrite dynamics, we examined animals that developed entirely under light, in which high-order branches clustered near the neuronal soma. To visualize the dynamics of dendrites over a 50-min period, we mapped the positions of all dendrite tips in each frame (with an interval of 3 min) and projected all time points into a single image. Tip positions were color-coded so that growing and shrinking tips can be distinguished. Our analysis shows that terminal branches of WT neurons are highly dynamic throughout the arbor (Figure 8C and Movie S7), exhibiting branch elimination (closed arrowheads, indicated by the absence of warm colors), branch addition (open arrowheads, indicated by the absence of cold colors), and repeated retraction and extension (open arrows, indicated by overlapping tracks of cold and warm colors). In contrast, the dendrites of Khc-trapped neurons were much more static (Figure 8D and Movie S8), with most tips exhibiting slow retraction (closed arrows, indicated by distal cold and proximal warm colors). Slow extension (open arrows) was observed only at a few branches very close to the soma.

Together, the above data suggest that kinesin-1 activity is required to support dynamic growth of dendrites.

### Kinesin-1 is required for ER delivery to high order branches

Rab5-mediated endosomes and Golgi outposts have been previously proposed to be the cargos of kinesin and dynein in supporting dendrite growth. Given that kinesin-1 transports a wide range of cargos, we wondered if other organelles could also be responsible for the dendrite defects caused by *Khc* LOF. We examined ER distribution in dendrites using the ER marker Sec61β-mCherry (Ferrandiz et al., 2022), because ER is the location for the synthesis of membrane and secretory proteins, the origin of the secretory pathway, and a cargo of kinesin (Lippincott-Schwartz et al., 2000). In WT neurons, ER was continuously distributed in both low- and high-order branches, even in proximal segments of terminal dendrites (Figures 8E and 8E’, closed arrowheads). In contrast, in Khc-trapped neurons (Figures 8F and 8F’), ER was almost entirely absent from high-order branches (open arrowheads) and exhibited gaps in low-order branches (arrows). As a result, ER occupied a much smaller portion of dendrites in these neurons as compared to the WT (Figure 8G). These results suggest that kinesin-1 is responsible for delivering ER to high-order, including terminal, dendrite branches. Considering that ER plays fundamental roles in many cellular processes, defects in ER delivery could be an important contributing factor to the dendrite phenotypes caused by *Khc* LOF.

## DISCUSSION

### OptoTrap for light control of endogenous proteins in *Drosophila*

In this study, we report a protein trapping system that allows manipulation of endogenous proteins in *Drosophila* using blue light. This system was designed to offer great versatility and superior spatiotemporal resolution for *in vivo* protein manipulation. We demonstrate the effectiveness of this system by trapping a range of endogenous proteins in epithelial cells and neurons.

The versatility of this system is expanded by choices at three levels: the prey/bait pair, pMag variants, and the copy number of nMag. First, we engineered two prey/bait pairs: NB/GFP and split GFP fragments. Because many endogenous proteins in *Drosophila* have already been tagged by GFP (Li-Kroeger et al., 2018; Lowe et al., 2014; Venken et al., 2011), the NB versions of OptoTrap can work with these existing, off-the-shelf reagents. On the other hand, the split GFP system offers several unique advantages: (1) Because GFP_11_ is very small (16 amino acids), tagging a protein with even several copies of GFP_11_ usually causes less disruption to protein function than tagging with the full GFP. Thus, GFP_11_-tagged strains are usually healthier, which can be important for generating homozygotes (in which all proteins of interest are tagged and thus can be trapped) containing OptoTrap and other necessary components. (2) With GFP_11_ as the tag, the POI can be selectively labeled in specific tissues by expressing GFP_1-10_ only in those tissues (Kamiyama et al., 2016). (3) Tagging the POI with multiple copies of GFP_11_ can increase the trapping efficiency due to crosslinking among CRY2olig clusters.

Second, we incorporate two variants of pMag that show different association and dissociation kinetics. In practice, these variants show distinct recovery time and allow for manipulation at different temporal scales. The fast variant provides precise temporal and spatial control and thus is suitable for manipulating signaling events with high temporal precision. In contrast, the slow variant is continuously active for hours with a single irradiation, making it ideal for long-term trapping of the POI.

Lastly, we designed 1X- and 2X-nMag versions for use in different tissues. Our results show that the two versions behave differently in neurons and epidermal cells, likely due to the distinct geometries of the two cell types. Epidermal cells represent cells with simple shapes, whereas neurons have long and slender branches. While 1X-nMag is effective in epidermal cells, 2X-nMag is needed in the limited volume of neuronal branches, likely due to enhanced aggregation by crosslinking.

With OptoTrap, we demonstrated rapid protein trapping at subcellular resolution, such as in part of an epithelial cell and selected branches of a neuron. Designed to manipulate endogenous proteins in model organisms, this system is distinct from most other optogenetic tools developed to date in two ways. First, most previous tools rely on overexpression of exogenous designer proteins that produce specific signaling outputs (Johnson and Toettcher, 2019; Shao et al., 2018; Toettcher et al., 2013; Wu et al., 2009; Zhao et al., 2019). Although these tools are useful for inducing artificial dominant active effects, they are ineffective in revealing physiological functions of proteins. In contrast, OptoTrap acts directly on endogenous proteins and can reveal their spatiotemporal requirements in specific cell types. Second, previous approaches of developing light-controllable agents typically require protein-specific, labor-intensive optimization and hence is difficult to apply to broader proteins. In comparison, OptoTrap can be easily applied to a wide range of proteins that are tagged for multiple purposes. Thus, OptoTrap offers a unique, powerful, and low-cost option for finely dissecting physiological functions of numerous genes.

### Practical factors in the design and applications of OptoTrap

While developing OptoTrap, we noticed two factors that are critical for the success of our strategy. First, as CRY2olig can form light-independent clusters at high expression levels, introduction of Magnets in the two-step recruitment design is necessary for minimizing the system’s dark activity: Even if CRY2olig forms clusters in the dark, recruitment of nMag-prey to the clusters still depends on light, ensuring precise control of the POI. Second, the relative expression levels of CRY2olig-pMag(3X) and nMag-prey have a significant impact on trapping. Theoretically, complete recruitment of nMag-prey to CRY2olig-pMag clusters requires overwhelmingly more pMag available for nMag binding. Including three copies of pMag in the CRY2olig-pMag moiety is not sufficient by itself, as our earlier designs of expressing the two parts at similar levels were not effective. Our solution is to express CRY2olig-pMag(3X) with a high-expression UAS vector and nMag-prey with a low-expression UAS vector (Han et al., 2011). Similarly, nMag-prey should be expressed at a much higher level than the POI, which is relatively easy to achieve as the Gal4/UAS-driven nMag-prey is usually expressed more highly than typical endogenous proteins.

When applying OptoTrap, additional considerations need to be made for the desirable outcome. Because the system operates by trapping or clustering proteins, it may affect the activity of different types of proteins differently. Theoretically, this system is most effective for inducing LOF of proteins whose activity relies on being at certain subcellular location, such as motors mediating cargo transport. However, for proteins whose activity is naturally regulated by the state of clustering, OptoTrap could induce GOF rather than LOF. This latter property could be utilized to understand how protein activity is regulated. For example, our results of optogenetic manipulation of Nrg supports the idea of Nrg activation by clustering (Jefford and Dubreuil, 2000).

Lastly, at least three control experiments are necessary for accurate assessment of the results. First, expression of OptoTrap in WT cells under light is needed to ensure that aggregation of the system is neutral in the cell type of interest. Second, nMag-prey expression in homozygotes of the tagged gene can reveal whether binding of nMag-prey to the POI affects its function. Lastly, expression of OptoTrap in homozygotes of the tagged gene in the dark should not produce strong phenotype and should serve as the baseline for light manipulation.

### OptoTrap reveals roles of MT in dendrite maintenance

Although MTs are known to be important for neuronal morphogenesis (Iwanski and Kapitein, 2023; Kelliher et al., 2019), how MTs support the growth of highly dynamic dendritic branches is unclear. Because it is difficult to detect MTs in dynamic terminal dendrite branches, investigating the local role of MTs in dynamic dendrites has been challenging. Traditional LOF approaches lack the spatial and temporal resolution needed to fully address this question. Using OptoTrap, we demonstrate that MTs are required for the maintenance of high-order branches of C4da neurons. In particular, acute trapping of α-tubulin causes immediate retraction of terminal branches. OptoTrap could affect MTs in these branches by sequestering free, tagged tubulin monomers/dimers and/or by directly disrupting dynamic MT filaments that have incorporated tagged *α*-tubulin. However, we think reduction of tubulin monomers/dimers (50% at the maximum) alone cannot completely explain the results, as early αTub84B-GFP_11_ trapping (which should reduce *α*-tubulin concentration and also prevent incorporation of αTub84B-GFP_11_ into dynamic MT filaments) did not affect high-order branches. Regardless of the exact mechanism, these data are consistent with the idea that MTs do exist in terminal branches and support branch growth by mediating cargo transport and/or by providing mechanical support. The presence of ER in WT terminal dendrites, which is dependent on kinesin-1, further indicates that MT-mediated cargo transport occurs in terminal dendrites. Because markers for stable and bundled MTs cannot be detected in those branches (Poe et al., 2017), MTs are more likely individual and dynamic filaments that are easily disrupted by OptoTrap.

### OptoTrap reveals spatiotemporal requirements of kinesin-1 in dendrite patterning

MTs contribute to neuronal morphogenesis by supporting motor-based cargo transport, which is demonstrated by the striking dendrite phenotypes of *khc* and *dynein* mutant neurons (Satoh et al., 2008; Zheng et al., 2008). However, how motors contribute to dendrite growth at different stages of neuronal differentiation could not be addressed by conventional methods. Using OptoTrap to perturb Khc in different temporal windows of C4da dendrite growth, we discovered that kinesin-1 affects the final patterns of low- and high-order branches differentially in a temporal-specific manners: For the growth of low-order branches, which correlates with the arbor size, kinesin-1 is more important in the first 72 h; in contrast, the last 48 h of neuronal development is the critical window for the growth of high-order branches. In addition, while 48-72 h are minimally required for reactivated kinesin-1 to rescue growth of low-order branches, 24-48 h of kinesin-1 recovery is sufficient for reactivating the growth of high-order branches. A possible explanation for these differences is that the growth of low- and high-order branches requires different cargos that are transported at unequal rates. Because kinesin-1 is mostly kept in the cell body with prolonged trapping and it takes less than 30 min to relieve kinesin-1 inhibition, delivery of critical cargos from the cell body may take >48 h for low-order branches and >24 h for high order branches.

Our time-lapse imaging experiments provided additional clues on how kinesin-1 supports dendrite growth. We found that kinesin-1 is required for dynamic growth of dendrites. Increasing length of inhibition results in faster retraction of dynamic branches, likely due to more severe local depletion of growth-promoting cargos. Prolonged kinesin-1 inhibition further leads to mostly static or slowly retracting dendrite tips. Mechanistically, in addition to the previously reported cargos Rab5-positive endosomes and Golgi outposts (Kelliher et al., 2018; Satoh et al., 2008; Zheng et al., 2008), we found that ER is a cargo that requires kinesin-1 for delivery to terminal dendrites and for its integrity in low-order branches. Because ER is involved in fundamental cellular activities ranging from protein synthesis to membrane trafficking, the ER abnormalities caused by kinesin-1 trapping could contribute significantly to dendrite defects.

Furthermore, our experiments reveal a temporal requirement of kinesin-1 in affecting the spatial distribution of high-order branches. Although proximal shift of high-order branches is a hallmark of kinesin-1 mutant clones (Satoh et al., 2008), to produce this phenotype, both persistent suppression in the last 96 h and early activity in the first 24 h are necessary. Counterintuitively, additional suppression of kinesin-1 in the first 24 h (as in the 120 h light group) results in weaker proximal shift and milder dendrite reduction. A possible explanation is that complete suppression of kinesin-1 in newly born neurons enhances the activities of other kinesins that could partially compensate for the loss of kinesin-1, while kinesin-1 activity in the first 24 h suppresses such compensatory mechanisms. Following this reasoning, *Khc* mutant C4da clones in otherwise heterozygous animals can generate consistent dendrite proximal shift (Satoh et al., 2008) possibly due to residual wild-type Khc proteins inherited from neuronal progenitor cells at the time of clone generation. Two other observations are also consistent with possible activities of other kinesins. First, even in compete suppression of kinesin-1, ER can still be delivered into low-order branches. Second, even in the worst phenotype of kinesin-1 LOF, neurons are still able to grow dendritic arbors spanning more than 200 microns. These phenomena can unlikely be explained by diffusion alone.

Lastly, our comparison of animals derived from homozygous and heterozygous mothers showed that post-mitotic neurons can inherit maternally contributed kinesin-1 and that this presumably residual amount of kinesin-1 can impact dendrite patterns. Because OptoTrap can access all endogenous proteins in a cell, if they are all tagged as illustrated in this example, to our knowledge, OptoTrap is the only known method that can eliminate the effects of maternal contribution and perdurance at single-cell level.

### Possible applications of OptoTrap in other systems

In principle, OptoTrap can be applied to other tissues in *Drosophila*, if those tissues can be penetrated by blue light. We found that OptoTrap can even be activated in body wall tissues of freely growing larvae in normal media under ambient light while displaying very little dark activity. The tight control, versatility, and the cell-type specificity of the system makes it a potentially powerful tool for dissecting many developmental processes *in vivo*. Being modular, the prey-bait pair in the system can be replaced to accommodate endogenous proteins containing other tags. OptoTrap may also be applied to other model organisms that are amenable to transgenic expression and light access.

## METHODS

The details of fly strains used in this study are listed in Key Resource Table. For expression in epidermal cells, we use *Gal4^R15A11^*as an intermediate driver, *Gal4^R16D01^* and *Gal4^en^*as strong drivers that are expressed in strips of epidermal cells. For expression in C4da neurons, we used *Gal4^ppk^*, except in Figures 7 and S7 where we used *Gal4^21-7^*, because *Khc-GFP_11x7_* homozygotes with *ppk-Gal4* were lethal.

See Supplemental Methods for details of molecular cloning and transgenic flies, live imaging, photactivation, immunostaining, image analysis and quantification, and statistical analysis.

## Supporting information

Key resource table

## ACKNOWLEDGEMENTS

We thank Dr. Lingfeng Tang, Dr. Hui Ji, and Yue Qiu for testing earlier versions of OptoTrap; Diana Bank and Ziqing Zhong for writing Dendrite Arbor Analyzer plugin; Dr. Jean-Yves Tinevez for writing the Dendrite Dynamic Tracker plugin; Addgene for plasmids; Bloomington Stock Center for fly stocks; the Cornell Statistical Consulting Unit (CSCU) for aiding statistics analysis; and Dr. Andrew Thomas Lombardo for discussion and suggestions on the manuscript. This work was supported by NIH grants (R21OD023824, R01NS099125, and R24OD031953) awarded to C.H., and by NIH grant (R01NS102385) awarded to J.W.

## DECLARATION OF INTEREST

The authors declare no competing interests.

---

**Figure S1.**
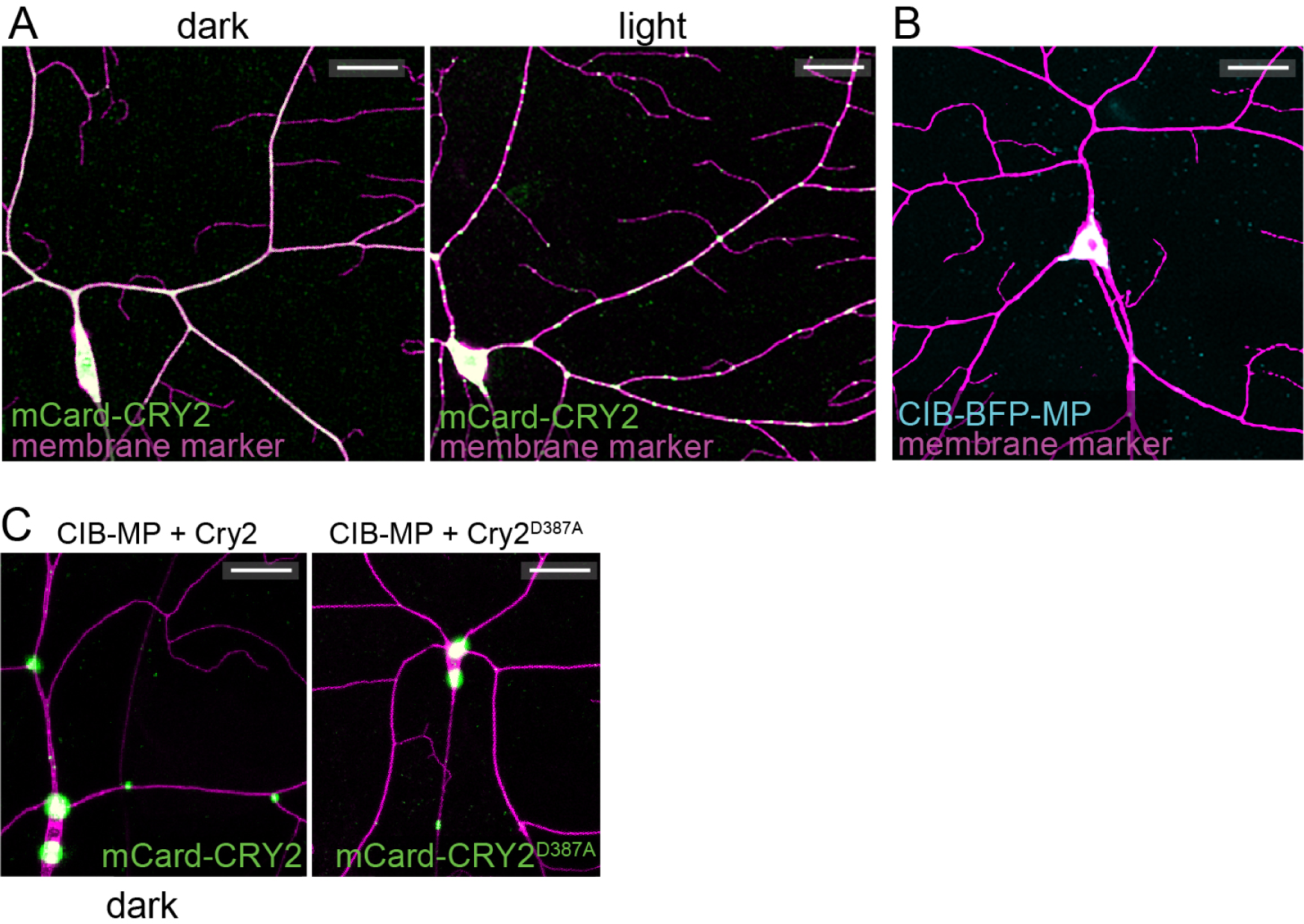
The LARIAT system is not compatible with *Drosophila* C4da neurons. (A) C4da neurons in *Gal4^ppk^>mCard-CRY2* in the dark and light. (B) A C4da neuron in *Gal4^ppk^>CIB-BFP-MP*. (C) C4da neurons in *Gal4^ppk^>CIB-MP + CRY2* in the dark and *Gal4^ppk^>CIB-MP + CRY2^D387A^*. In all panels, neurons were labeled by CD4-tdTom; scale bars represent 50 μm.

**Figure S2.**
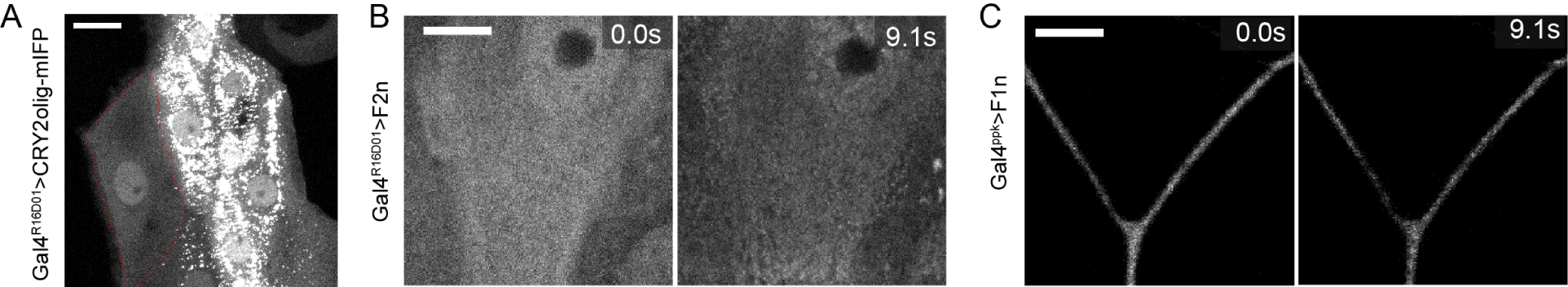
Characterization of OptoTrap. (A) Epidermal cells in *Gal4^R16D01^>CRY2olig-mIFP* in the dark. Scale bars, 25 μm (B) Epidermal cells in Gal4^R16D01^>OptoTrap (F2n) animals before (0.0 s) and after (9.1 s) blue laser activation. Scale bar, 10 μm (C) A C4da neuron in *Gal4^ppk^>OptoTrap^F1n^* before (0.0 s) and after (9.1 s) blue laser activation. Scale bar, 10 μm

**Figure S3.**
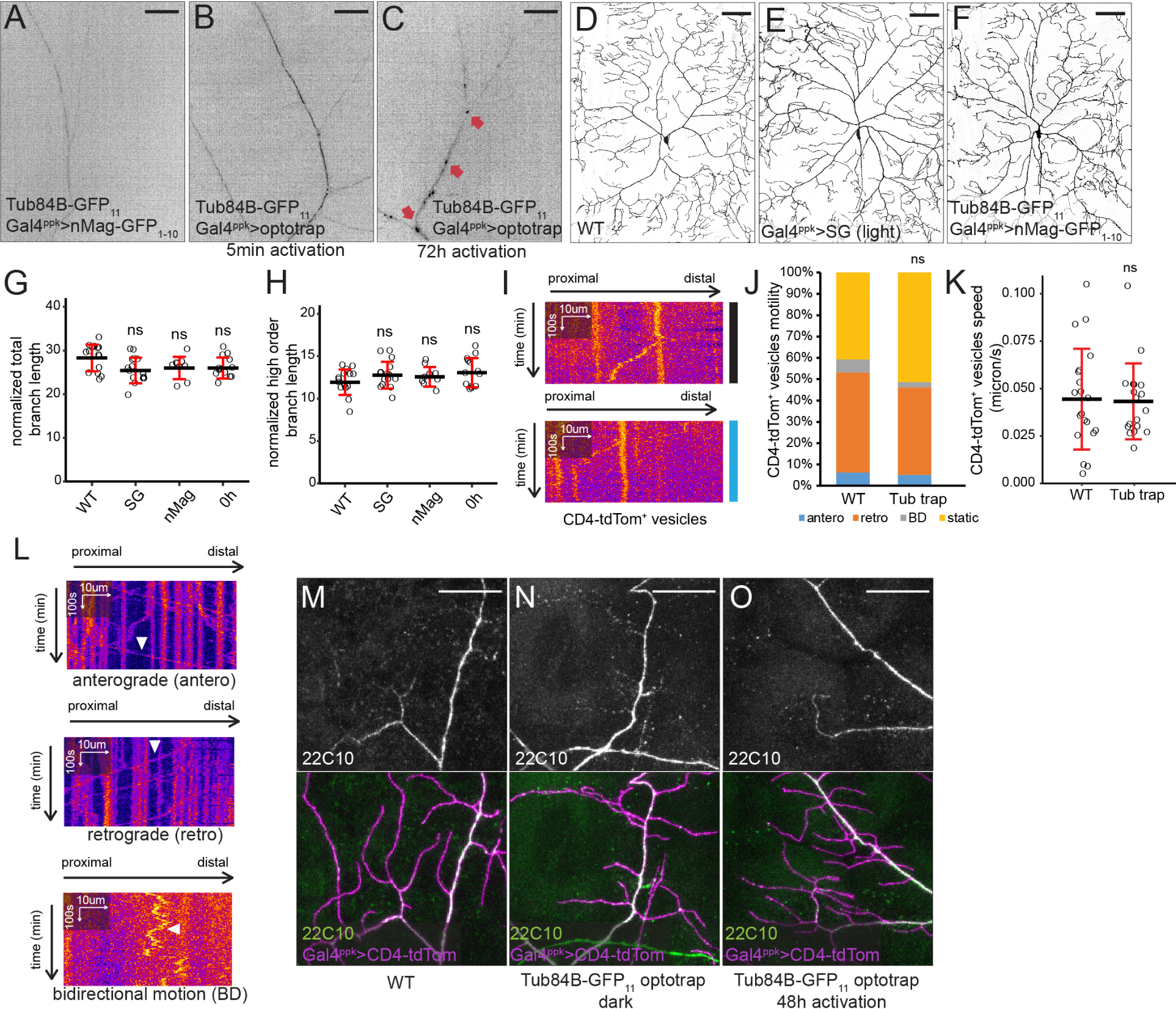
Effects of *α*-Tub84B-GFP_11_ trapping. (A-C) GFP signals reconstituted by α-Tub84B-GFP_11_ and nMag-GFP_1-10_ in C4da neurons in *Gal4^ppk^>nMag-GFP_1-10_* (A), *Gal4^ppk^>OptoTrap^SG^* that were under light for 5 min before imaging (B), and *Gal4^ppk^>OptoTrap^SG^* that were reared under light for 72h before imaging (C). Images were taken by structured illumination microscopy (SIM). Scale bars, 20 μm. (D-F) C4da neurons labeled by *Gal4^ppk^>CD4-tdTom* in WT (D), *Gal4^ppk^>OptoTrap^SG^* reared under light (E), *α-Tub84B-GFP_11_*/+; *Gal4^ppk^>nMag-GFP_1-10_* (F). Scale bars, 100 μm (G-H) Total branch length (G) and high-order branch length (H) normalized by segment width. Each circle represents a neuron. n=15 for WT, n=15 for OptoTrap^SG^ under light (SG); n=9 for nMag-GFP_11_ (nMag); n=13 for *Gal4^ppk^>OptoTrap^SG^; α-Tub84B-GFP_11_/+* in the dark (0 h). ns, not significant; One-way ANOVA and Tukey’s HSD test. Black bars, mean; red bars, SD. The 0 h data are the same as in Figure 5G and 5H. (I) Kymographs of CD4-tdTom-labeled vesicles in C4da neurons of *Gal4^ppk^>OptoTrap^SG^; α-Tub84B-GFP_11_/+* that grew in the dark (top) or exposed to light from the beginning of imaging (bottom). (J) Motility of CD4-tdTom-labeled vesicles. n=32 for WT; n=39 for acute trapping of α-Tub84B-GFP_11_ (Tub trap). ns, not significant; Freeman–Halton extension of Fisher’s exact test. (K) Speed of CD4-tdTom-labeled vesicles among the moving (anterograde, retrograde and bidirectional motion) populations in (J). Each circle represents a vesicle. n=20 for WT; n=18 for Tub trap. ns, not significant; One-way ANOVA and Tukey’s HSD test. Black bars, mean; red bars, SD. (L) Example kymographs to show anterograde, retrograde and bidirectional motion vesicles. (M-O) Anti-Futsch (22C10) staining in WT (M) and *Gal4^ppk^>OptoTrap^SG^; α-Tub84B-GFP_11_/+* animals that grew in the dark (N) or were reared under light for 48 h before imaging (M). Dendrites were labeled by CD4-tdTom. Scale bars, 10 μm.

**Figure S4.**
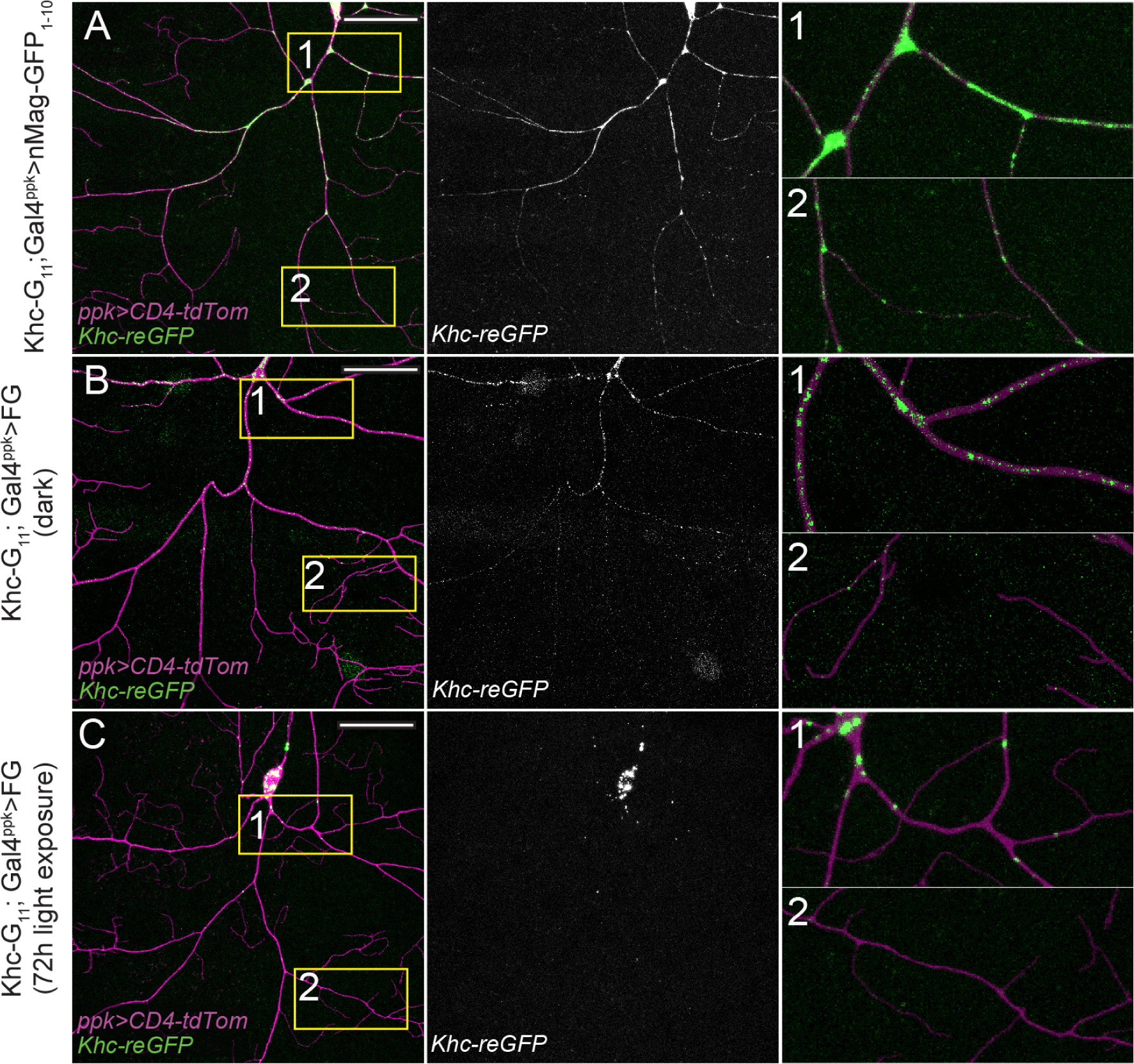
OptoTrap restricts Kinesin distribution in C4da neurons. (A-C) C4da neurons in Khc-GFP_11x7_ homozygotes expressing *Gal4^ppk^>nMag-GFP_1-10_* (A), *Gal4^ppk^>OptoTrap^FG^* that were reared in the dark (B), *Gal4^ppk^>OptoTrap^FG^* that were reared under light for 72h before imaging. Imagining reconstituted GFP (Khc-reGFP) exposes the neuron to blue laser. (1) proximal region. (2) distal region. Scale bars, 50 μm

**Figure S5.**
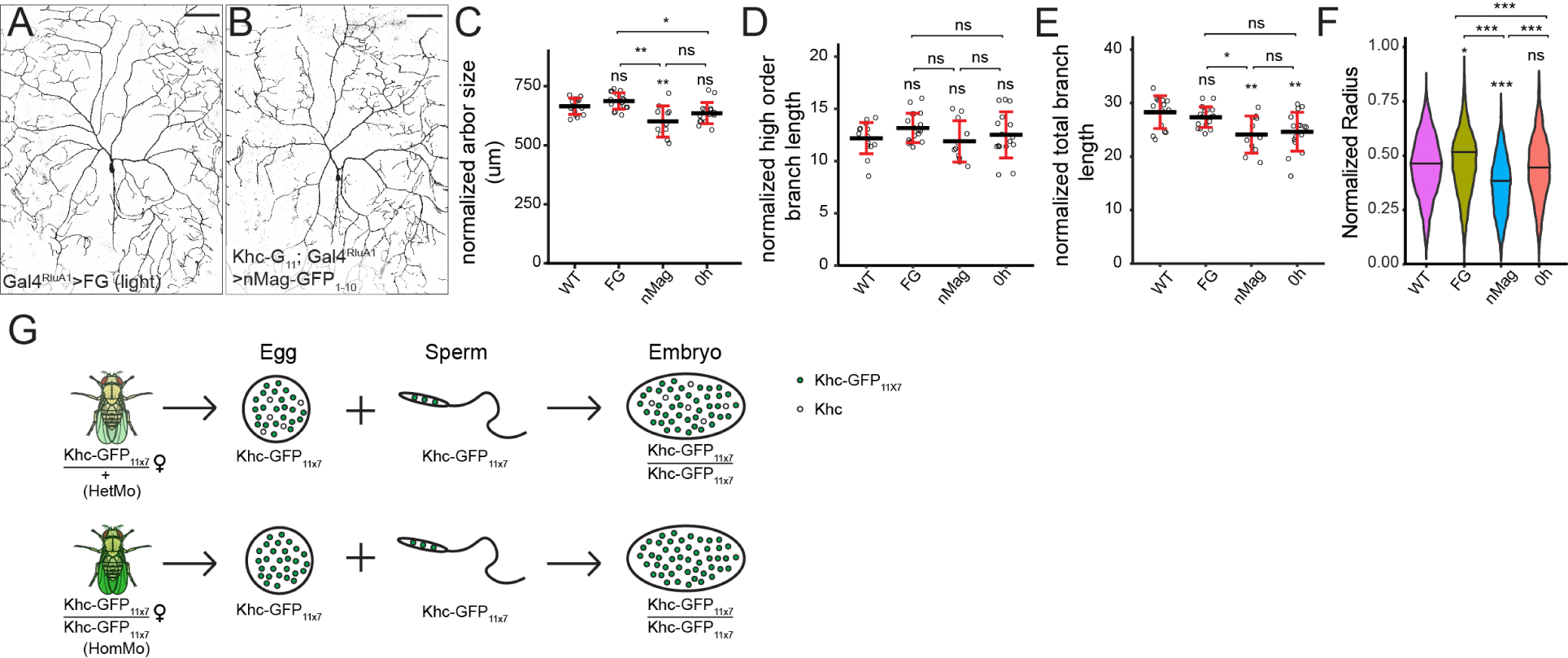
OptoTrap controls in C4da neurons. (A and B) C4da neurons in *Gal4^RluA1^>OptoTrap^FG^* that grew under light (A) and *Khc-GFP_11x7_; Gal4^RluA1^>nMag-GFP_1-10_* (B). (C-E) Arbor size (C), high-order branch length (D), and total branch length (E) of C4da neurons normalized by segment width. Each circle represents a neuron. (F) Sholl analysis of high-order branches of C4da neurons presented as violin plots. In (D-F), n=15 for WT, n=18 for *Gal4^RluA1^>OptoTrap^FG^*that grew under light (FG); n=11 for *Khc-GFP_11x7_; Gal4^RluA1^>nMag-GFP_1-10_* (nMag); n=16 for 0 h in light. The dataset of 0 h in light is the same as 120 h dark in Figure 7. (G) A diagram showing the difference between *Khc-GFP11_X7_* homozygous and heterozygous mothers. HetMo: heterozygous mother; HomMo: homozygous mother. Green circles represent Khc-GFP11_X7_; white circles represent untagged Khc. For all quantifications, \*\**p*<0.01; \**p*<0.05; ns, not significant; One-way ANOVA and Tukey’s HSD test. Black bars, mean; red bars, SD.

**Table S1.**
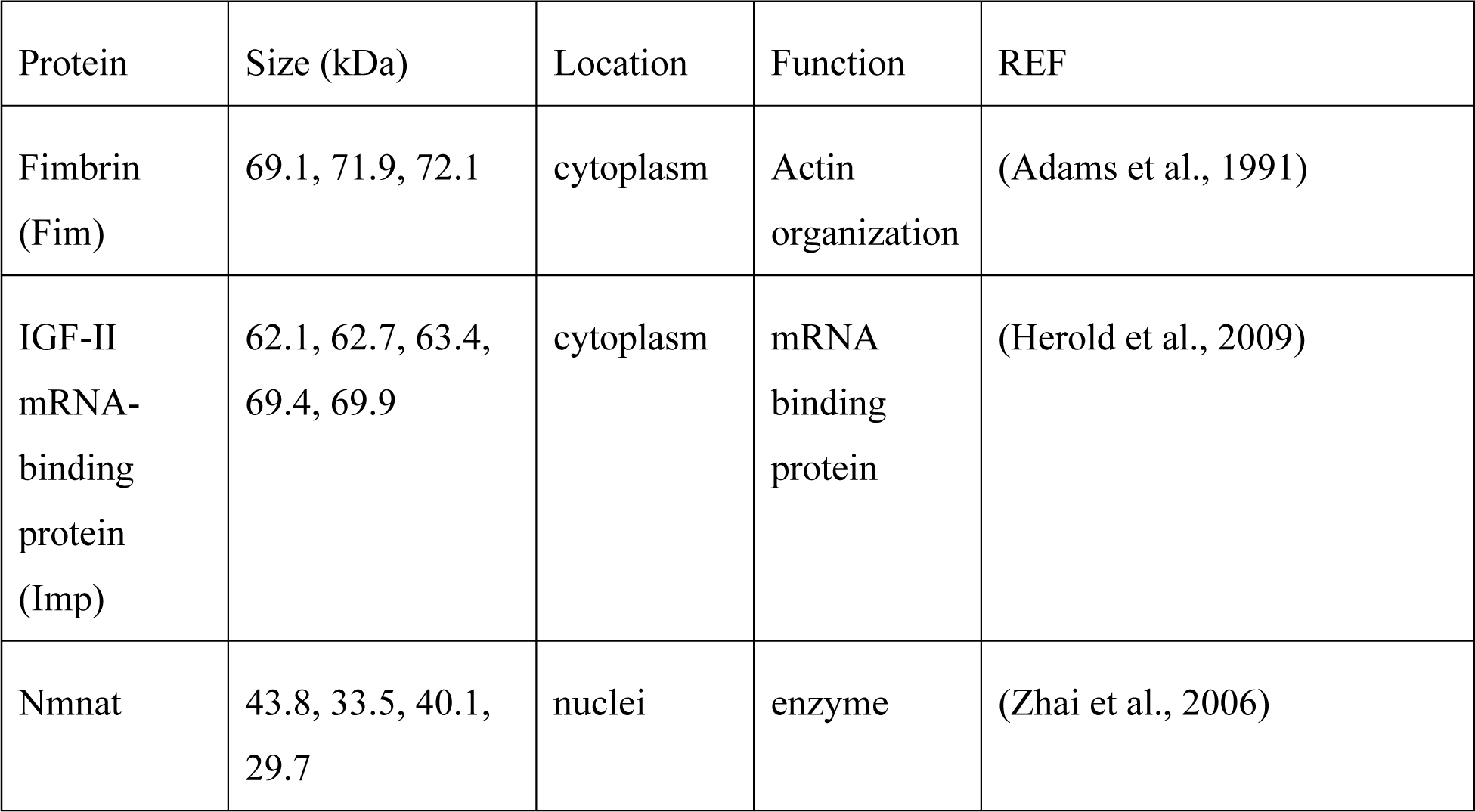
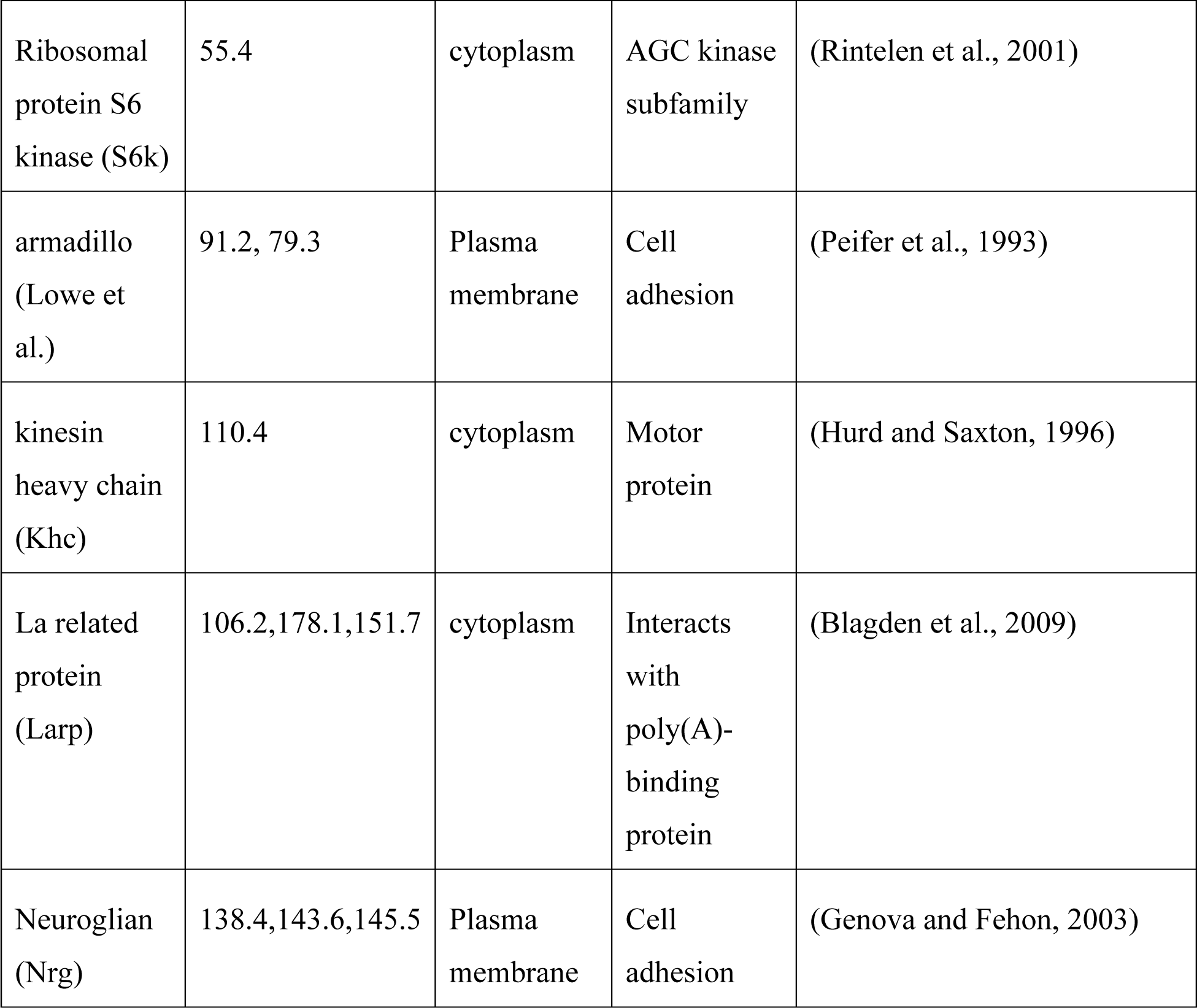

| Protein | Size (kDa) | Location | Function | REF |
| --- | --- | --- | --- | --- |
| Fimbrin<br>(Fim) | 69.1, 71.9, 72.1 | cytoplasm | Actin<br>organization | (Adams et al., 1991) |
| IGF-II<br>mRNA-binding<br>protein<br>(Imp) | 62.1, 62.7, 63.4,<br>69.4, 69.9 | cytoplasm | mRNA<br>binding<br>protein | (Herold et al., 2009) |
| Nmnat | 43.8, 33.5, 40.1,<br>29.7 | nuclei | enzyme | (Zhai et al., 2006) |
| Ribosomal protein S6 kinase (S6k) | 55.4 | cytoplasm | AGC kinase subfamily | (Rintelen et al., 2001) |
| armadillo (Lowe et al.) | 91.2, 79.3 | Plasma membrane | Cell adhesion | (Peifer et al., 1993) |
| kinesin heavy chain (Khc) | 110.4 | cytoplasm | Motor protein | (Hurd and Saxton, 1996) |
| La related protein (Larp) | 106.2,178.1,151.7 | cytoplasm | Interacts with poly(A)-binding protein | (Blagden et al., 2009) |
| Neuroglian (Nrg) | 138.4,143.6,145.5 | Plasma membrane | Cell adhesion | (Genova and Fehon, 2003) |

## MOVIE LEGENDS

**Movie S1. Local activation of OptoTrap in epidermal cells.**

Larval epidermal cells in *Gal4^R16D01^>OptoTrap^S1n^*. The OptoTrap was visualized by mIFP. The animal was reared in the dark. Only the boxed region was illuminated by 488 nm laser from the beginning of the movie.

**Movie S2. Local activation of OptoTrap in neurons.**

A C4da neuron in *Gal4^ppk^>OptoTrap^F2n^*. OptoTrap was visualized by mIFP, with *UAS-HO1* co-expressed to enable mIFP fluorescence. The animal was reared in the dark. Only the dendrite segment in the boxed region was illuminated by 488 nm laser from the beginning of the movie.

**Movie S3-S5. Microtubules are important for dendrite branch dynamics.**

C4da neurons in the WT (S3), α-Tub84B 48 h trapping (S4) and α-Tub84B acute trapping (S5) in *Gal4^ppk^>OptoTrap^SG^; α-Tub84B-GFP_11_/+* animals. Neurons were labeled by *Gal4^ppk^>CD4-tdTom*.

**Movie S6. Local trapping of Khc inhibits mitochondrial transport in neurons.**

A C4da neuron in a *Khc-GFP_11x7_* homozygous larva that expresses *Gal4^ppk^>OptoTrap^FG^* and the mitochondrial marker mito-mCherry. The animal was reared in the dark. Only the boxed region was illuminated by 488 nm laser from the beginning of the movie; the rest of the neuron remained in the dark.

**Movie S7-S8. WT and Khc-trapped neurons exhibit different branch dynamics.**

C4da neurons in the WT (S7) and a *Khc-GFP_11x7_; Gal4^ppk^>OptoTrap^SG^* HetMo larva that was reared in constant light (S8). Neurons were labeled by *Gal4^ppk^>CD4-tdTom.* Dendrite endings in each frame were marked by dots in a color that corresponds to the temporal sequence of the frame.

## Notes

### Competing Interest Statement

The authors have declared no competing interest.

