## Supplementary material for "Light-induced trapping of endogenous proteins reveals spatiotemporal roles of microtubule and kinesin-1 in dendrite patterning of *Drosophila* sensory neurons": Key resource table

| Reagent type | Designation in the text | Source or reference | identifiers | Additional information |
|---|---|---|---|---|
| Fly strain | *UAS-mCard-Cry2* | This study | | *UAS-mCard-Cry2*$^{VK37}$ |
| Fly strain | *UAS-CIB-BFP-MP* | This study | | *UAS-CIB-TagBFP2-MP*$^{VK16}$ |
| Fly strain | *UAS-mCard-Cry2*$^{D387A}$ | This study | | *UAS-mCard-Cry2(D387A)* $^{VK37}$ |
| Fly strain | *Nrg-GFP* | Bloomington Drosophila Stock Center | RRID: BDSC_6844 | *Nrg*$^{G00305}$ |
| Fly strain | *R16D01-Gal4* | Bloomington Drosophila Stock Center | RRID: BDSC_48722 | *R16D01-Gal4*$^{attP2}$ |
| Fly strain | *ppk-Gal4* | (Han et al., 2012) | | *ppk-Gal4*$^{1a}$ |
| Fly strain | *UAS-CD4-tdTom* | (Han et al., 2011) | RRID: BDSC_35841 | *UAS-CD4-tdTom*$^{7M1}$ |
| Fly strain | *UAS- mIFP-2A-HO1* | (Poe et al., 2017) | RRID: BDSC_64181 | *UAS-mIFP-T2A-HO1*$^{VK00005}$ |
| Fly strain | *R15A11-Gal4* | Bloomington Drosophila Stock Center | RRID: BDSC_48674 | pruned by BDSC |
| Fly strain | *en-Gal4* | (Han et al, 2014) | | |
| Fly strain | *RluA1-Gal4* | This study | | |
| Fly strain | *ppk-CD4-tdTom* | (Han et al., 2012) | | *ppk-spGFP11-CD4-tdTom*$^{2}$ |
| Fly strain | *Fim-GFP* | Bloomington Drosophila Stock Center | RRID: BDSC_59838 | |
| Fly strain | *Larp-GFP* | Bloomington Drosophila Stock Center | RRID: BDSC_61790 | |
| Fly strain | *Imp-GFP* | Bloomington Drosophila Stock Center | RRID: BDSC_60237 | |
| Fly strain | *Khc-GFP* | (Winding et al., 2016) | | |
| Fly strain | *Nmnat-GFP* | Bloomington Drosophila Stock Center | RRID: BDSC_80087 | |
| Fly strain | *S6k-GFP* | Bloomington Drosophila Stock Center | RRID: BDSC_51563 | |

| | | | | |
|---|---|---|---|---|
| Fly strain | *Khc-GFP$_{11}$* | (Winding et al., 2016) | | |
| Fly strain | *α-Tub84B-GFP$_{11}$* | This study | | |
| Fly strain | *UAS-Nrg* | Bloomington Drosophila Stock Center | RRID: BDSC_64181 | |
| Fly strain | *UAS-Nrg-RNAi* | Bloomington Drosophila Stock Center | RRID: BDSC_64181 | |
| Fly strain | *UAS-mito-mCherry* | Bloomington Drosophila Stock Center | RRID: BDSC_66533 | |
| Fly strain | *UAS-Sec61β-mCherry* | Bloomington Drosophila Stock Center | RRID: BDSC_64746 | |
| Fly strain | *UAS-HO1* | This study | | |
| Fly strain | *UAS-Cry2olig-mIFP* | This study | | |
| Fly strain | *UAS- Cry2olig-pMagFast2(3x)* | This study | | |
| Fly strain | *UAS- Cry2olig-pMag(3x)* | This study | | |
| Fly strain | *UAS-mIFP-nMagHigh1-NB* | This study | | |
| Fly strain | *UAS-mIFP-nMagHigh1(2x)-NB* | This study | | |
| Fly strain | *UAS-mIFP-nMagHigh1-GFP1-10* | This study | | |
| Antibody | Rat anti-D-CAD2 | Developmental Studies Hybridoma Bank | | 1:200 dilution |
| Antibody | Donkey Anti-Rat Cy3 | Jackson ImmunoResearch Laboratories | RRID: AB_2340667 | 1:400 dilution |
| Antibody | Mouse anti-Futsch | Developmental Studies Hybridoma Bank | | 1:500 dilution |
| Antibody | Donkey Anti-mouse Cy3 | Jackson ImmunoResearch Laboratories | RRID: AB_ 2315777 | 1:400 dilution |
| Software | FIJI | https://fiji.sc/ | RRID: SCR_002285 | |

| | | | | |
|---|---|---|---|---|
| Software | R | https://www.r-project.org/ | RRID: SCR_001905 | |
| Food ingredient | Agar | MoorAgar, Inc | Cat #: 41084 | Gelidium |
| Food ingredient | Glucose | LD Carlson Company | Cat #: 1992 | Dextrose-corn sugar |
| Food ingredient | Inactive yeast | VWR International, LLC | Cat #: IC90331280 | Yeast (brewers), Powder |
| Chemical | Phosphoric Acid | VWR International, LLC | Cat #: JT0260-2 | |
| Chemical | Propionic Acid | Krackeler Scientific | Cat #: 45-P1386-1L | |
